# 4q-D4Z4 chromatin architecture regulates the transcription of muscle atrophic genes in FSHD

**DOI:** 10.1101/623363

**Authors:** Alice Cortesi, Matthieu Pesant, Shruti Sinha, Federica Marasca, Eleonora Sala, Francesco Gregoretti, Laura Antonelli, Gennaro Oliva, Chiara Chiereghin, Giulia Soldà, Beatrice Bodega

## Abstract

Despite increasing insights in genome structure organization, the role of DNA repetitive elements, accounting for more than two thirds of the human genome, remains elusive. Facioscapulohumeral Dystrophy (FSHD) is associated with deletion of D4Z4 repeat array below 11 units at 4q35.2. It is known that the deletion alters chromatin structure *in cis*, leading to genes upregulation. Here we show a genome-wide role of 4q-D4Z4 array in modulating gene expression via 3D nuclear contacts. We have developed an integrated strategy of 4q-D4Z4 specific 4C-seq and chromatin segmentation analyses, showing that 4q-D4Z4 3D interactome and chromatin states of interacting genes are impaired in FSHD1 condition; in particular, genes which have lost the 4q-D4Z4 interaction and with a more active chromatin state are enriched for muscle atrophy transcriptional signature. Expression level of these genes is restored by the interaction with an ectopic 4q-D4Z4 array, suggesting that the repeat directly modulates the transcription of contacted targets.

Of note, the upregulation of atrophic genes is a common feature of several FSHD1 and FSHD2 patients, indicating that we have identified a core set of deregulated genes involved in FSHD pathophysiology.

## Introduction

Among primate specific macrosatellites, D4Z4 is a 3.3 Kb unit tandem repeat duplicated on several chromosomes (Bakker et al. 1995; Deidda et al. 1995; Lyle et al. 1995; Bodega et al. 2006; Bodega et al. 2007), and in particular present as a polymorphic array of 11 to 100-150 copies at 4q35.2 (4q-D4Z4 array) in the general population (Hewitt et al. 1994). Reduction of 4q-D4Z4 array copy number below 11 units is associated with Facioscapulohumeral Dystrophy (FSHD, MIM158900; (van Deutekom et al. 1993)), one of the most common myopathies in humans with an overall prevalence of more than 1:10,000 (Sacconi et al. 2015). FSHD is characterized by progressive, often asymmetric, weakness and wasting of facial (facio), shoulder and upper arm (scapulohumeral) muscles (Tawil and Van Der Maarel 2006), where fiber necrosis and degeneration give rise to muscle atrophy (Sacconi et al. 2015).

FSHD is a genetically variable disorder, mainly transmitted as an autosomal dominant trait, on a specific FSHD-permissive haplotype of Chromosome 4q, namely 4qA (Lemmers et al. 2002; Lemmers et al. 2007). This form accounts for approximately 95% of the cases (FSHD1); however, about 5% of the patients display FSHD lacking D4Z4 array contractions (FSHD2). FSHD2 is caused by mutations in *SMCHD1*, a member of the condensin/cohesin chromatin compaction complexes, that binds to the D4Z4 repeat array (Lemmers et al. 2012). While in healthy individuals the 4q-D4Z4 array is characterized by highly methylated DNA, the contracted allele in FSHD1 and both the 4q-D4Z4 alleles in FSHD2 are hypomethylated (van Overveld et al. 2003; de Greef et al. 2009).

The highly heterogeneous FSHD clinical features suggest a strong epigenetic contribution to the pathology (Tawil et al. 1993; Cabianca and Gabellini 2010; Neguembor and Gabellini 2010; Lanzuolo 2012; Daxinger et al. 2015). It is described that the 4q-D4Z4 array is able to engage short- and long-range genomic contacts with several genes *in cis* (Petrov et al. 2006; Bodega et al. 2009; Himeda et al. 2014; Robin et al. 2015), concomitantly to the Polycomb group (PcG) protein binding and histone deacetylation, resulting in an overall chromatin compaction. Instead, in FSHD1 condition such interactions are lost, with the consequent alteration of the chromatin structure at the FSHD locus, leading to a more active chromatin state, which is responsible for the de-repression of the genes *in cis* (Gabellini et al. 2002; Jiang et al. 2003; Bodega et al. 2009; Zeng et al. 2009; Cabianca et al. 2012). In particular, one of the major player in FSHD pathogenesis is the transcription factor *DUX4*, encoded from the most telomeric D4Z4 repeat (Gabriels et al. 1999; Dixit et al. 2007; Lemmers et al. 2010); DUX4 is normally silenced in somatic cells (Snider et al. 2010), but it has been found overexpressed in FSHD patients’ myotubes, leading to the activation of genes associated with RNA metabolism processes, stem cell and germ-line development, MERVL/HERVL retrotransposons (Geng et al. 2012; Young et al. 2013; Rickard et al. 2015; Hendrickson et al. 2017) and resulting in the induction of toxicity and apoptosis of muscle cells (Bosnakovski et al. 2008; Block et al. 2013).

Besides the established role of 4q-D4Z4 array in modulating the transcription of *in cis* genes, whether the repeat could also directly affect chromatin structure and gene expression of other loci via 3D physical contacts has not been investigated yet. Therefore, we have explored the 4q-D4Z4 chromatin architecture and possible alterations in FSHD.

## Results

### 4q-specific D4Z4 interactome is deregulated in FSHD1 patients

Given the high duplication and sequence similarity of 4q35.2 with multiple regions of the genome, in particular with 10q26.3 (Bodega et al. 2006; Bodega et al. 2007), we designed a 4q-specific 4C-seq (circular chromosome conformation capture sequencing) strategy to investigate its interactome. As 4C viewpoint (VP) we used the region nearby a single sequence length polymorphism (SSLP), present shortly upstream (almost 3.5 Kb) of the first D4Z4 repeat on 4q (4qA and 4qB) and 10q arrays (Lemmers et al. 2007); we performed paired-end sequencing, that allowed to retrieve the information of the SSLP variant (Read 1) and the interacting region (Read 2), assigning with high precision the allele origin of the D4Z4 interactome (Supplemental Fig. S1; Supplemental Table S1; see Methods and Supplemental Methods).

With this approach, we probed the 4q-D4Z4 chromatin conformation in human primary muscle cells from two FSHD1 patients (FSHD1) and two healthy individuals (CN) (Supplemental Table S1), that did not differ for myoblast purity and differentiation efficiency (Supplemental Fig. S2). 4C-seq was performed on myoblasts (MB) to highlight differences that could precede any transcriptional effect in differentiated cells. Comparative analyses of 4C-seq samples showed high level of reproducibility and similarity both at the level of donor origin (fragends read count, CN or FSHD1) (Supplemental Fig. S3) and at the level of viewpoint (called interacting regions, 4q vs 10q) (Supplemental Fig. S4A,B). We identified 4q-D4Z4 specific *cis* interactions with *FRG1, ZFP42, SORBS2* and *FAM149A* genes (Fig. 1A; Supplemental Fig. S4C-E), as already reported (Bodega et al. 2009; Robin et al. 2015), suggesting that our approach is robust in the detection of 4q-specific D4Z4 interactions.

**Figure 1.**
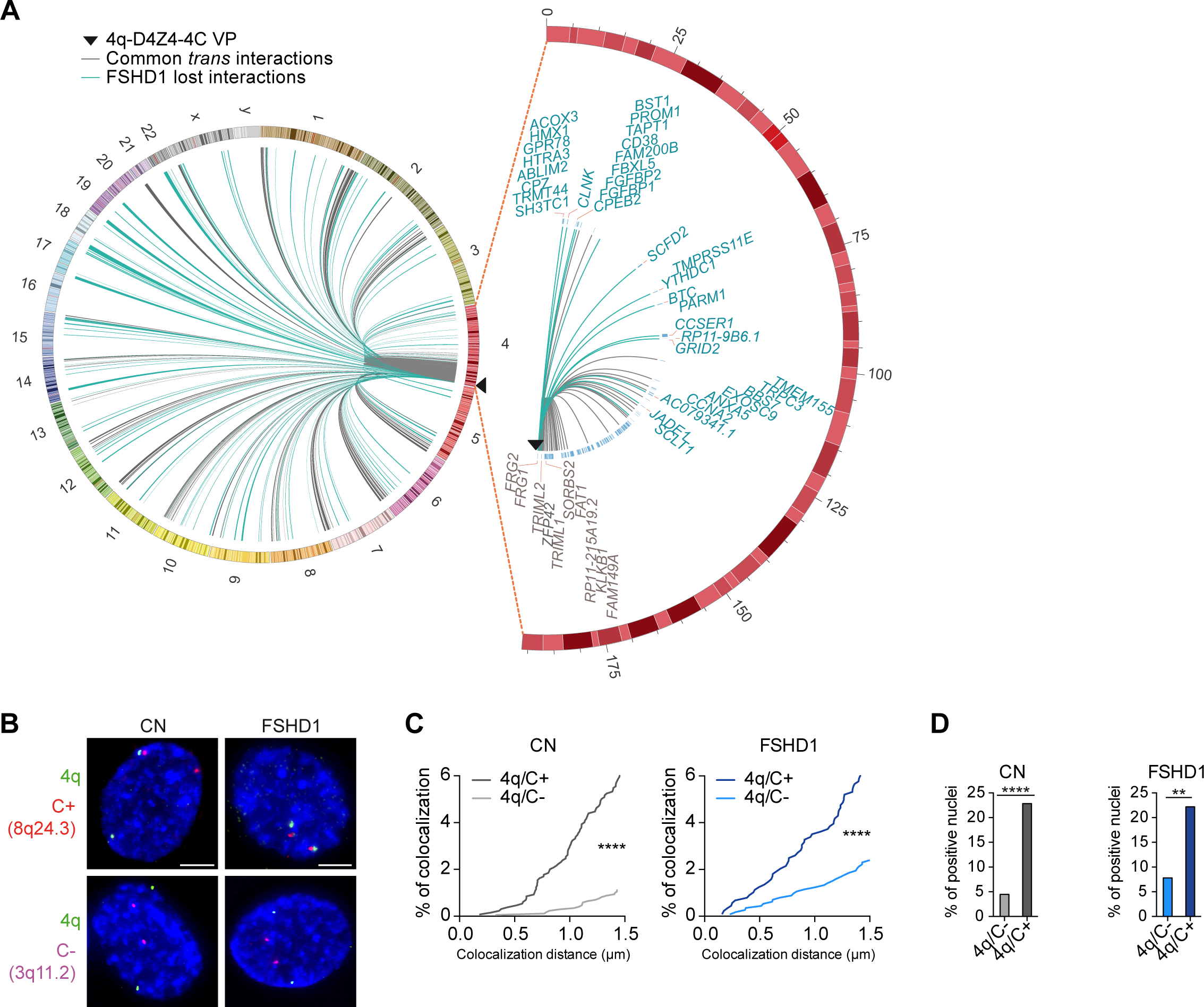
4q-D4Z4 specific 4C-seq highlights FSHD1 impaired interactome. (*A*) (left) Circos plot depicting *cis* and *trans* 4q-D4Z4 interactions in CN (CN-3, CN-4) myoblasts called by 4C-ker. Common interactions with FSHD1 (FSHD1-3, FSHD1-4) myoblasts are in grey, whereas interactions specifically lost in FSHD1 are highlighted in light blue. (right) Zoomed-in circos plot representation of common (grey) and FSHD1 lost (light blue) *cis* interactions on Chr 4. Gene are indicated for a region extending up to 4 Mb from the VP. Black triangles in circos plots depict the VP localization. (*B*) Representative nuclei of 3D multicolor DNA FISH using probes mapping to 4q35.1 region (4q, green), a 4q-D4Z4 positive interacting region (8q24.3, C+, red) and a 4q-D4Z4 not interacting region (3q11.2, C-, magenta) and in CN (CN-1, CN-2, CN-3, CN-4) and FSHD1 (FSHD1-1, FSHD1-2, FSHD1-3, FSHD1-4) myoblasts. Nuclei are counterstained with DAPI (blue). All images at 63X magnification. Scale bar=5 µm. (*C*) Cumulative frequency distributions of distances (below 1.5 µm) between 4q and C+ and between 4q and C-in CN (dark and light grey; left) and FSHD1 (dark and light blue; right) myoblasts. n=1,296 (CN 4q/C+), 1,708 (CN 4q/C-), 884 (FSHD1 4q/C+) and 1,128 (FSHD1 4q/C-). *P* values were calculated by unpaired one-tailed *t*-test with confidence interval of 99%. Asterisks represent statistical *P* values; for 4q/C+ vs 4q/C-in CN and FSHD1 *p*<0.0001. (*D*) Percentage of nuclei positive for the interactions (under the cut-off of 1.5 µm). n= 427 (CN 4q/C-), 324 (CN 4q/C+), 282 (FSHD1 4q/C-) and 221 (FSHD1 4q/C+). *P* values were calculated by fisher’s exact one-sided test with confidence interval of 99%. Asterisks represent statistical *P* values; for 4q/C-vs. 4q/C+ in CN *p*<0.0001; for 4q/C-vs. 4q/C+ in FSHD1 *p*=0.0046.

We retrieved 244 and 258 4q-D4Z4 interacting regions for CN and FSHD1 respectively, and in particular, among them, 175 for CN and 181 for FSHD1 were *trans* interactions. Interestingly, 116 regions interacting in CN were specifically lost in FSHD1 cells and the vast majority (101) were in *trans* (Fig. 1A; Supplemental Table S2).

3D multicolor DNA FISH was performed on the same and additional CN and FSHD1 donor MB to validate 4C results, using a probe on a not duplicated region in 4q35.1 (Supplemental Fig. S5A; (Tam et al. 2004)), and a probe for a positive (C+) or negative (C-) 4q-D4Z4 interacting region (Fig. 1B; Supplemental Fig. S5B). We developed a novel algorithm (NuCLεD, Nuclear Contacts Locator in 3D, see Supplemental Methods) to automatically detect and localize fluorescent spots in 3D reconstructed nuclei. We observed that 4q/C+ interaction had higher frequency of contacts and higher number of positive interacting nuclei compared to 4q/C-interaction (Fig. 1C,D; Supplemental Table S3), with contact frequencies in the range of those estimated for long range interactions (10-20%, (Finn et al. 2019)). Furthermore, 4q and C+ regions shared the same topological nuclear domain in both CN and FSHD1, whereas 4q and C-did not (Supplemental Fig. S5C,D; Supplemental Table S3). Same results were obtained in CN and FSHD1 myotubes (MT) (Supplemental Fig. S5E-G; Supplemental Table S3). Additionally, with our 4C-seq approach we were also able to retrieve 4q allele specific interactomes (4qA and 4qB), as well as 10q-D4Z4 interactome (Supplemental Fig. S6; Supplemental Table S2; see Supplemental Material).

Overall, the 4q-D4Z4-4C-seq strategy allowed to map genome-wide 4q-D4Z4 contacts and to highlight those deregulated in FSHD1.

### Genes that show impaired 4q-D4Z4 interactions and activated chromatin state are enriched for atrophic transcriptional signature

In order to identify novel deregulated genes specific for the FSHD condition, we derived chromatin state changes in FSHD1 cells and intersected with 4q-D4Z4 lost interactions in FSHD1, retrieving genes altered both at structural and chromatin levels.

To define the chromatin state, we generated or used available (ENCODE) ChIP-seq datasets for H3K36me3, H3K4me1, H3K27ac, H3K4me3 and H3K27me3 in CN and FSHD1 MB and MT (Supplemental Fig. S7A,B). The quality of ChIP-seq was validated on the same and additional CN and FSHD1 donors MB and MT (Supplemental Fig. S7C-H). Next, we identified 15 chromatin states using ChromHMM (Ernst and Kellis 2012), that were adopted for further downstream analyses (Fig. 2A; Supplemental Fig. S8A,B). Interestingly, chromatin segmentation analysis revealed transitions at enhancers and promoters distinctive for FSHD1 cells (Supplemental Fig. S8C; see Supplemental Material).

**Figure 2.**
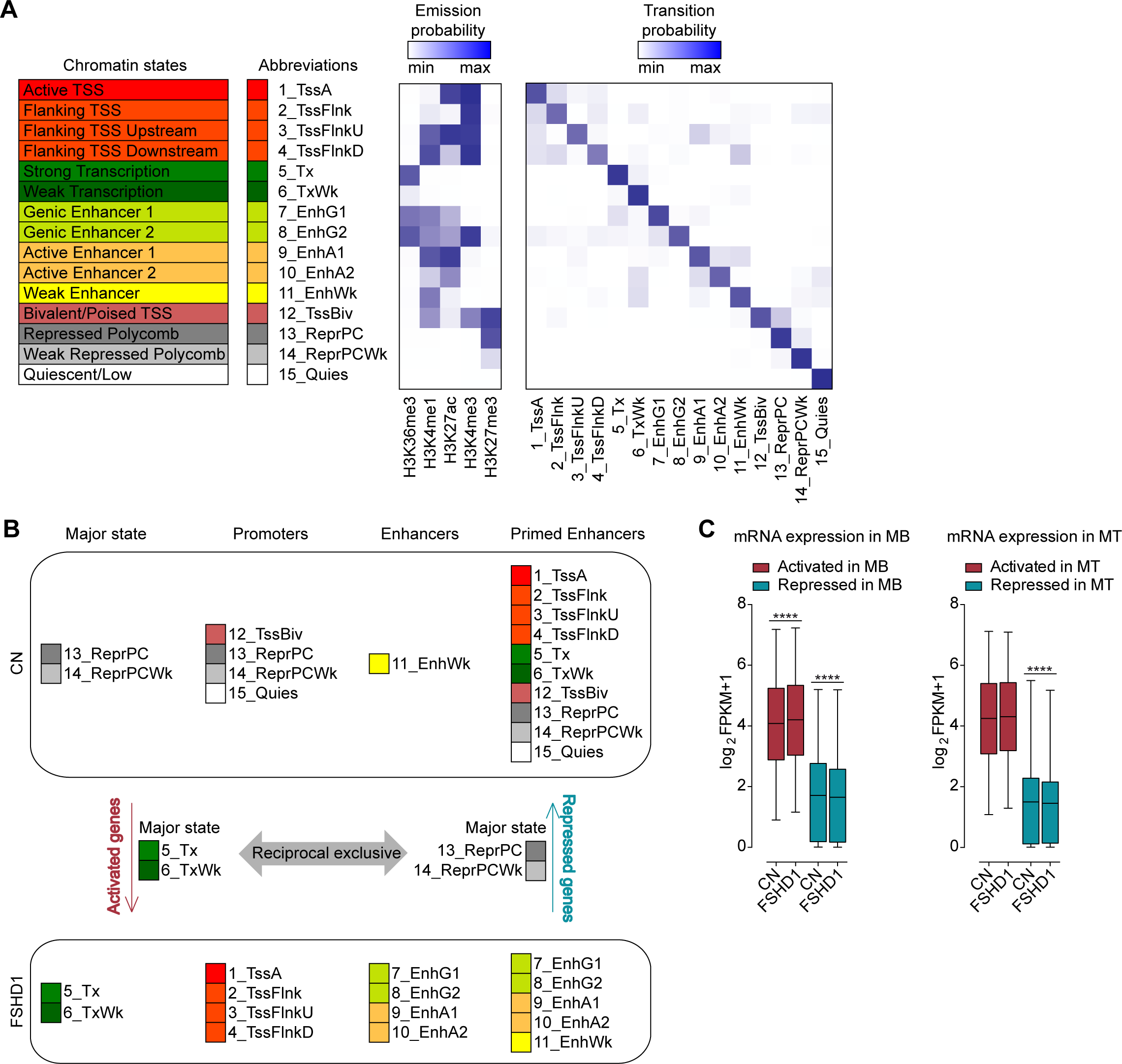
Chromatin segmentation analysis revealed chromatin state switches consistent with transcriptional changes in FSHD1 muscle cells. (*A*) ChromHMM 15-state model obtained with ChIP-seq datasets for H3K36me3, H3K4me1, H3K27ac, H3K4me3 and H3K27me3. Heatmaps display histone marks emission probabilities and transition probabilities between chromatin states. (*B*) Schematic representation of the strategy used to assign genes as activated or repressed in FSHD1. (*C*) Expression levels from RNA-seq datasets for FSHD1 activated and repressed genes in MB (left) and MT (right), in CN (CN-3, CN-4 and Yao’s datasets C20, C21, C22) and FSHD1 (FSHD1-3, FSHD1-4 and Yao’s datasets F4, F6) (Yao et al. 2014). Box & whiskers plots show the median of matched expression values of each gene for CN and FSHD1 and whiskers extend to the 5-95 percentiles. *P* values were calculated by paired two-tailed Wilcoxon matched-pairs signed rank test with confidence interval of 99%. Asterisks represent statistical *P* values; for CN vs. FSHD1 activated in MB *p*<0.0001; for CN vs. FSHD1 repressed in MB *p*<0.0001; for CN vs. FSHD1 repressed in MT *p*<0.0001.

In order to identify the genes that specifically switched to activated or repressed chromatin state in FSHD1, we designed the strategy shown in Fig. 2B. Activated genes were defined as those that showed a transition towards a more active state (considering the coverage of the gene body, promoter and enhancer regions) and repressed genes those that showed an opposite change (Fig. 2B; Supplemental Table S4; see Methods). To verify the reliability of this approach, we inspected the expression level of these genes by analyzing RNA-seq datasets performed on the corresponding cell lines and additional publicly-available RNA-seq datasets (Yao et al. 2014). Notably, the activated or repressed chromatin state switches were associated with higher or lower mRNA expression levels, respectively, in FSHD1 compared to CN (Fig. 2C; Supplemental Table S4).

We next sought genes that had lost the interaction with 4q-D4Z4 and also showed chromatin deregulation in FSHD1. We observed that 28% (450/1614) of genes that had lost contact with 4q-D4Z4 in FSHD1 were mainly activated (FSHD1 lost-activated genes, 71%, 319/450), whereas a minority of them (FSHD1 lost-repressed genes, 29%, 131/450) were repressed (Fig. 3A; Supplemental Fig. S9A-C; Supplemental Table S5; see Supplemental Material). Interestingly, only few of these FSHD1 altered genes were regulated by DUX4 (Supplemental Fig. S9D; Supplemental Fig. S10; see Supplemental Material).

**Figure 3.**
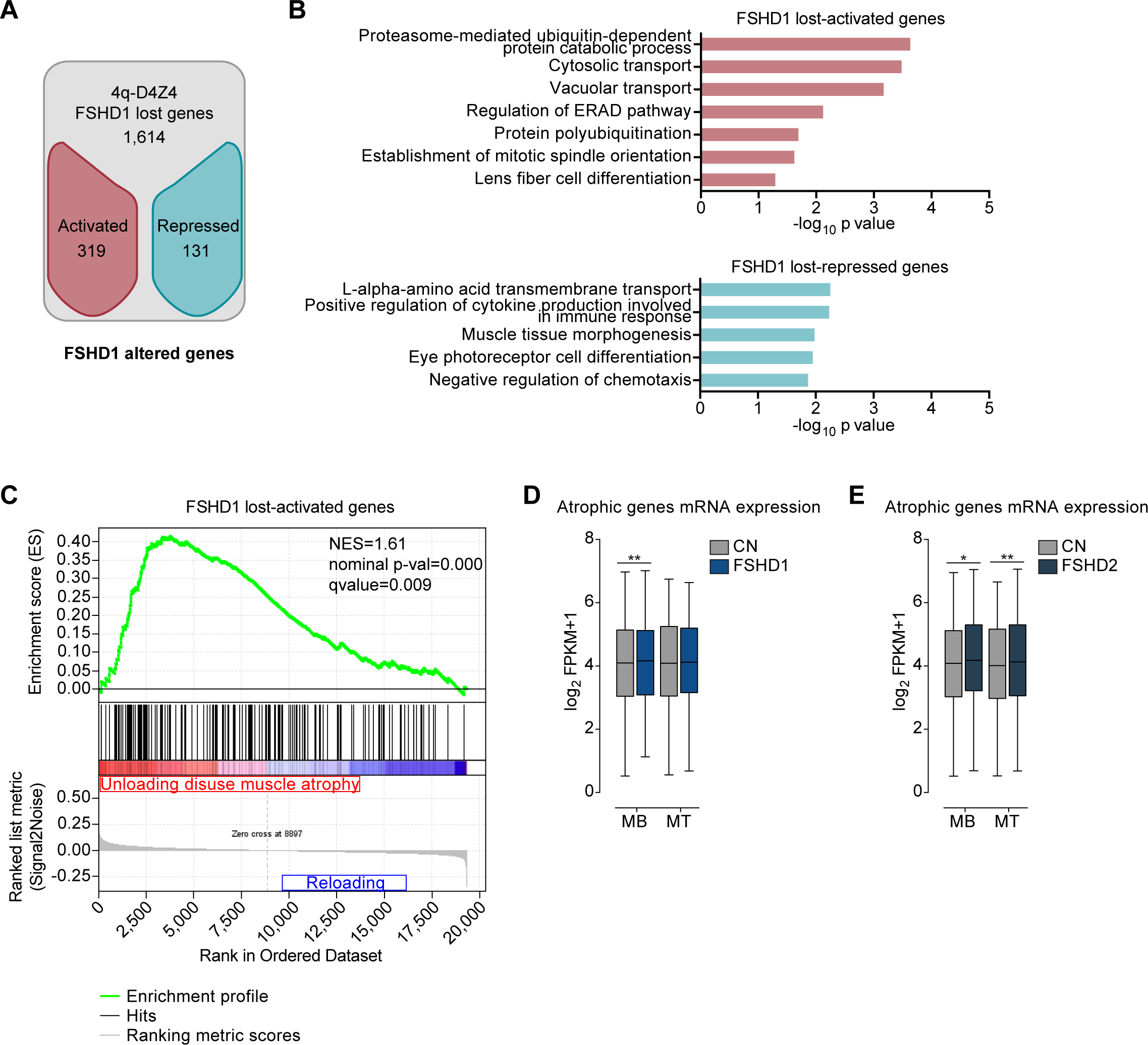
Genes which have lost the interaction with 4q-D4Z4 have a more active chromatin state and are enriched for muscle atrophy signature in FSHD muscle cells. (*A*) Flowchart of filtering steps to identify FSHD1 altered genes. Genes within lost 4q-D4Z4 interactions were filtered as activated (red) or repressed (blue) in FSHD1. (*B*) Gene Ontology analysis (Biological Processes) of FSHD1 lost-activated and repressed genes. Bars correspond to −log_10_ of the *P* value. (*C*) Gene Set Enrichment Analysis (GSEA) results of the 319 FSHD1 lost-activated genes performed on expression data from unloading-induced muscle atrophy subjects (Reich et al. 2010). Genes upregulated in atrophic condition are depicted in red whereas genes not enriched are depicted in blue. NES, Normalized Enrichment Score. (*D*) Expression levels from RNA-seq datasets for atrophic genes (Reich et al. 2010), in CN (CN-3, CN-4 and Yao’s datasets C20, C21, C22) and FSHD1 (FSHD1-3, FSHD1-4 and Yao’s datasets F4, F6) (Yao et al. 2014) MB and MT. Box & whiskers plots show the median of matched expression values of each gene for CN and FSHD1 and whiskers extend to the 5-95 percentiles. *P* values were calculated by paired two-tailed *t*-test with confidence interval of 99%. Asterisks represent statistical *P* values; for CN vs. FSHD1 in MB *p*=0.0099. (*E*) Expression levels from RNA-seq datasets for atrophic genes (Reich et al. 2010), in CN (CN-3, CN-4 and Yao’s datasets C20, C21, C22) and FSHD2 (Yao’s datasets F12, F14, F20) (Yao et al. 2014) MB and MT. Box & whiskers plots show the median of matched expression values of each gene for CN and FSHD2 and whiskers extend to the 5-95 percentiles. *P* values were calculated by paired two-tailed *t*-test with confidence interval of 99%. Asterisks represent statistical *P* values; for CN vs. FSHD2 in MB *p*=0.0251; for CN vs. FSHD2 in MT *p*=0.0041.

We performed Gene Ontology (GO) analyses on the FSHD1 altered genes and found that FSHD1 lost-activated genes were enriched in GO terms linked to protein catabolic processes and in particular with protein ubiquitination/degradation pathways (Fig. 3B; Supplemental Table S5), that are highly relevant to the FSHD-associated atrophic phenotype (Tawil and Van Der Maarel 2006; Sacconi et al. 2015; Statland and Tawil 2016). Similar analysis on 10q-D4Z4 FSHD1 altered genes did not reveal GO terms related to atrophy (Supplemental Fig. S11A,B).

Indeed, we executed Gene Set Enrichment Analysis (GSEA) and further demonstrated that 4q-D4Z4 specific lost-activated genes in FSHD1 were enriched for genes upregulated in the atrophic condition (Fig. 3C; Supplemental Fig. S11C-E; Supplemental Table S5). Of note, the FSHD1 lost-activated genes included in the atrophic dataset displayed higher expression level both in several FSHD1 (Fig. 3D) as well as FSHD2 (Fig. 3E) RNA-seq datasets, revealing that the epigenetic and transcriptional deregulation of this core set of genes represents a novel transcriptional signature that is common among different FSHD patients.

### 4q-D4Z4 lost interacting *FBXO32/ATROGIN1* gene is deregulated in FSHD patients at chromatin and transcriptional level

To finely dissect how the 4q-D4Z4 lost interactions could influence the upregulation of muscle atrophy genes in FSHD1 condition, we further investigated the regulation of *FBXO32* (*ATROGIN1*), that is one of the top-enriched genes identified by GSEA, and also one of the major player in different atrophy related conditions, in human and mouse (Gomes et al. 2001; Lecker et al. 2004; Sandri et al. 2004; Bodine and Baehr 2014). We verified by 3D multicolor DNA FISH the loss of *FBXO32*/4q-D4Z4 interaction on several FSHD1 donors compared to CN, extending our analysis also to FSHD2 (Fig. 4A; Supplemental Fig. S12A,B), and observed a decrease in *FBXO32*/4q-D4Z4 interaction frequency in FSHD1, but also in FSHD2 myoblasts (Fig. 4B,C; Supplemental Table S3). Similar results were obtained in FSHD1 myotubes, although with a smaller difference (Supplemental Fig. S12C-E; Supplemental Table S3).

**Figure 4.**
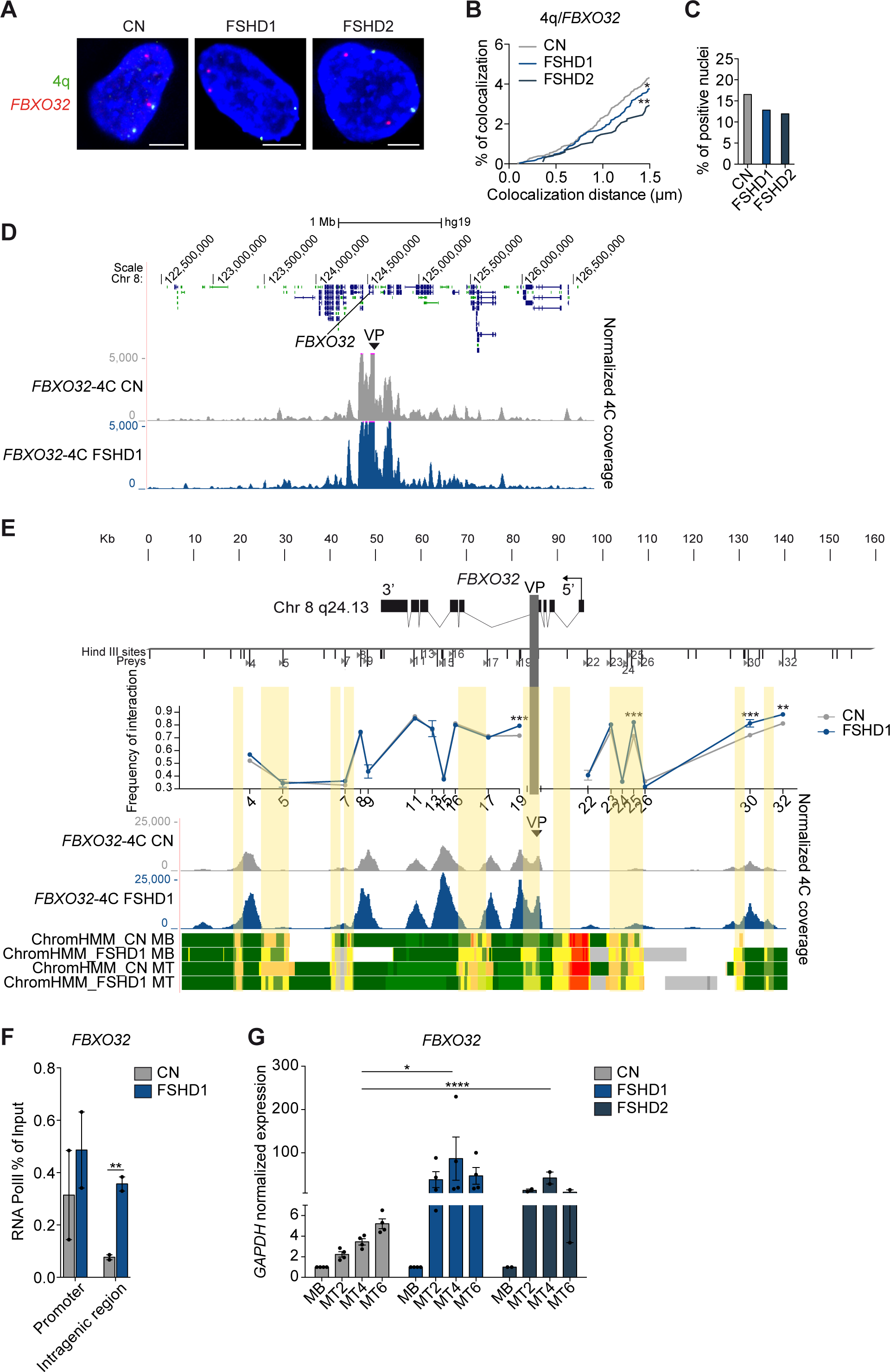
*FBXO32* gene has a deregulated chromatin structure and it is overexpressed in FSHD1 and FSHD2 muscle cells. (*A*) Representative nuclei of 3D multicolor DNA FISH using probes mapping to 4q35.1 region (4q, green) and *FBXO32* (red) in CN (CN-1, CN-3, CN-4), FSHD1 (FSHD1-1, FSHD1-3, FSHD1-4) and FSHD2 (FSHD2-1, FSHD2-2) myoblasts. Nuclei are counterstained with DAPI (blue). All images at 63X magnification. Scale bar=5 µm. (*B*) Cumulative frequency distribution of distances (below 1.5 µm) between 4q and *FBXO32* in CN (grey), FSHD1 (blue) and FSHD2 (dark blue) myoblasts. n= 3,652 (CN), 2,464 (FSHD1) and 1,020 (FSHD2). *P* values were calculated by unpaired one-tailed *t*-test with confidence interval of 99%. Asterisks represent statistical *P* values; for CN vs. FSHD1 *p*=0.0473; for CN vs. FSHD2 *p*=0.0036. (*C*) Percentage of nuclei positive for the interactions (under the cut-off of 1.5 µm). n= 913 (CN), 616 (FSHD1) and 255 (FSHD2). (*D*) 4C normalized coverage tracks at the *FBXO32* locus for *FBXO32*-4C VP in CN (CN-3, CN-4; grey) and FSHD1 (FSHD1-3, FSHD1-4; blue). (*E*) (top) Schematic representation of the *FBXO32* locus and Hind III sites. (middle) Chart showing the frequencies of 3C interaction between *FBXO32* promoter and the indicated Hind III restriction sites (sites 4-32), using the same bait of the 4C VP (light gray vertical bar) in CN (grey) and FSHD (blue). n=3 (CN) and 3 (FSHD1). S.e.m. is indicated. *P* values were calculated by two-way ANOVA followed by Bonferroni post-test correction. Asterisks represent statistical *P* values; for P19, P25 and P30 CN vs. FSHD1 *p*<0.001; for P32 CN vs. FSHD1 *p*<0.01. (bottom) 4C normalized coverage tracks as well as ChromHMM chromatin states tracks at the *FBXO32* locus for *FBXO32*-4C VP in CN (grey) and FSHD1 (blue). The arrow represents the promoter region; enhancers are highlighted in yellow. (*F*) Bar plot showing enrichment of RNA Pol II at *FBXO32* promoter (left) and an intragenic region (right) assessed by ChIP-qPCR experiment in CN (grey) and FSHD1 (blue) myoblasts. Results are presented as % of input. n=2 CN (CN-3, CN-4) and 2 FSHD1 (FSHD3, FSHD1-4). S.e.m. is indicated. *P* values were calculated by unpaired one-tailed *t*-test with confidence interval of 99%. Dots represent the values of each replicate; asterisks represent statistical *P* values; for *FBXO32* intragenic region CN vs. FSHD1 *p*=0.0050. (*G*) Expression levels of *FBXO32* gene during CN (grey), FSHD1 (blue) and FSHD2 (dark blue) differentiation (MB, myoblasts, MT2, myotubes day 2, MT4, myotubes day 4, MT6, myotubes day 6). Data were normalized on *GAPDH* expression and on MB. n=4 CN (CN-1, CN-2, CN-3, CN-4), 4 FSHD1 (FSHD1-1, FSHD1-2, FSHD1-3, FSHD1-4) and 2 FSHD2 (FSHD2-1, FSHD2-2). S.e.m. is indicated. *P* values were calculated by two-way ANOVA followed by Bonferroni post-test correction. Dots represent the values of each replicate; asterisks represent statistical *P* values; for MT4, CN vs. FSHD1 *p*<0.0290 and CN vs. FSHD2 *p*<0.0001.

*FBXO32* belongs to the category of activated genes in FSHD1, with the appearance of primed enhancers specifically in FSHD1 condition (Fig. 2B; Supplemental Fig. S13A; Supplemental Table S4). Therefore, we further investigated whether the *FBXO32* locus could display distinct chromatin loops at the level of enhancers-promoter in FSHD1. We performed 4C-seq (Fig. 4D; Supplemental Fig. S13B-D; Supplemental Table S1 and S2), showing interaction peaks between enhancers-promoter with higher normalized 4C reads coverage in FSHD1 (Fig. 4E;). These results were further corroborated in 3C experiments (Fig. 4E), suggesting a strengthening of enhancers-promoter contacts at *FBXO32* locus in FSHD1 cells.

In line with this observation, the binding of RNA Pol II at *FBXO32* promoter and an intragenic region was increased in FSHD1 myoblasts respect to CN (Fig. 4F; Supplemental Fig. S14A) and the *FBXO32* expression was upregulated in several FSHD1 donor muscle cells during differentiation, a trend that is also observed in FSHD2 (Fig. 4G). Finally, the *FBOX32* expression is not dependent by DUX4, as it is not affected by DUX4 overexpression (Supplemental Fig. S14B-D; see Supplemental Material), and ChIP-seq peaks (Geng et al. 2012) are absent in the *FBXO32* gene region (Supplemental Fig. S14E).

### Ectopic 4q-D4Z4 array restores the expression of FSHD1 lost interacting genes

To further investigate whether 4q-D4Z4 array could directly modulate the expression of interacting genes, we transfected CN and FSHD1 myoblasts with a BAC containing at least 15 D4Z4 repeat units (B Bodega, unpublished) (BAC 4q-D4Z4n from 4q35.2 region) in parallel with a control BAC (Ctrl BAC, unrelated and not interacting region); transfection efficiency was comparable among the BACs and ranging around 45% (Supplemental Fig. S14F-H; see Supplemental Methods).

We observed that specifically the ectopic 4q-D4Z4 array was in close spatial proximity to the endogenous 4q region, in 70% of the analyzed nuclei (Fig. 5A,B) and interacted with *FBXO32* with a frequency similar as that of the endogenous locus (roughly 20% of analyzed nuclei, Fig. 5C), indicating that the 4q-D4Z4 BAC occupies the same nuclear topological domain of the endogenous 4q region. We then assessed the effect of 4q-D4Z4 BAC transfection on the expression levels of a subset of genes that had lost 4q-D4Z4 interactions in FSHD1. We observed that the transcription of FSHD1 lost-activated genes was reduced (*FBXO32, TRIB3* and *ZNF555*; Fig. 5D), whereas a lost-repressed gene was upregulated (*LZTS3*; Fig. 5E) and no effect was detected for not interacting genes (*FOXO3 and MYOG*; Fig. 5F).

**Figure 5.**
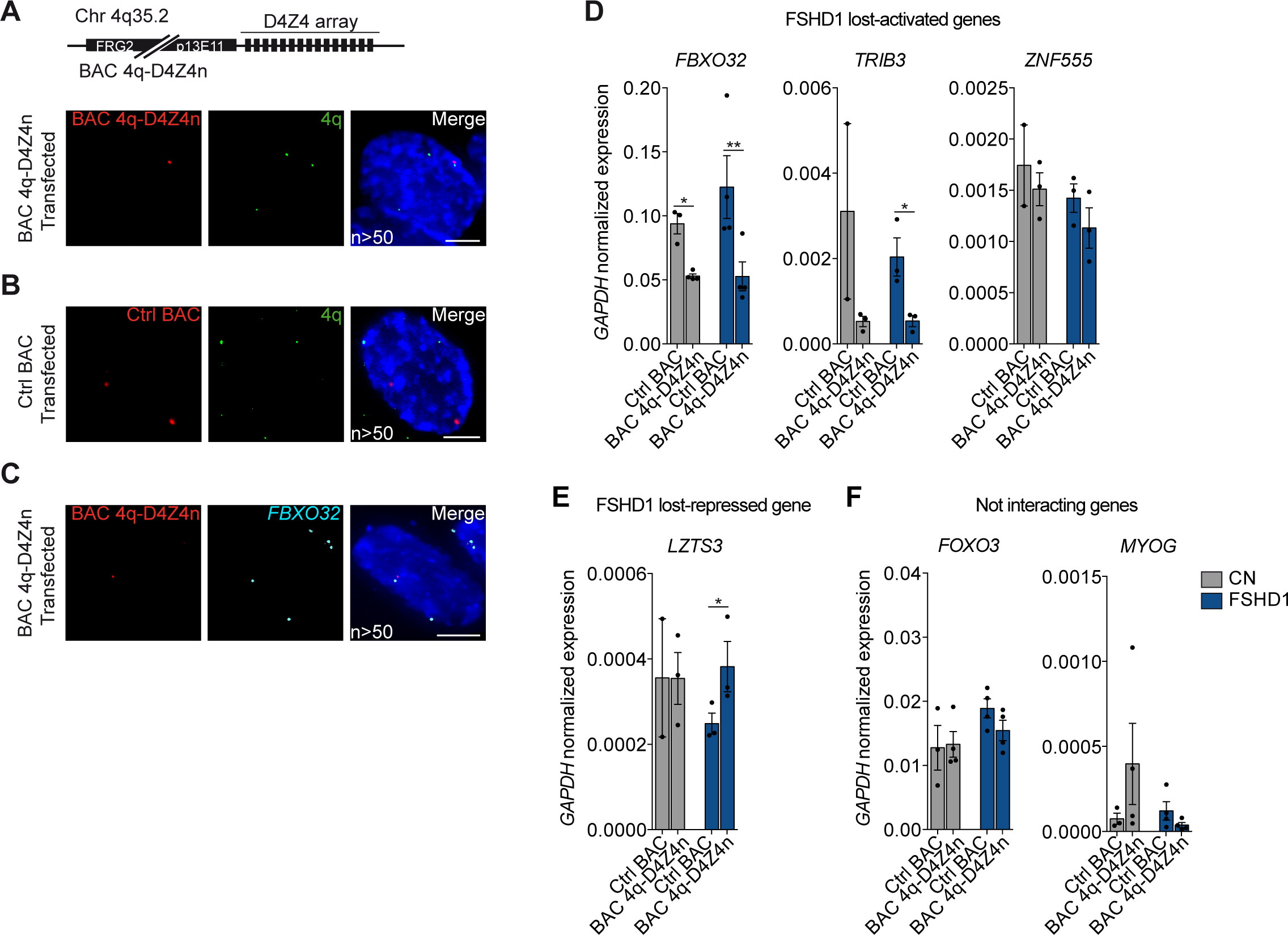
Ectopic 4q-D4Z4 array restores the expression of FSHD1 lost interacting genes. (*A*) (top) Representation of the BAC containing 4q upstream region and D4Z4 array (at least 15 D4Z4 units, B Bodega, unpublished (BAC 4q-D4Z4n)). (bottom) Representative nucleus of 3D multicolor DNA FISH using probes for the transfected BAC backbone (red) and 4q35.1 region (4q, green) in myoblasts transfected with BAC 4q-D4Z4n. Nuclei are counterstained with DAPI (blue). All images at 63X magnification. Scale bar=5 µm. n, number of nuclei analyzed. (*B*) Representative nucleus of 3D multicolor DNA FISH using probes for the transfected BAC backbone (red) and 4q35.1 region (4q, green) in myoblasts transfected with Ctrl BAC (RP11-2A16, representative of an unrelated and not interacting genomic region, Chr 17q21.33). Nuclei are counterstained with DAPI (blue). All images at 63X magnification. Scale bar=5 µm. n, number of nuclei analyzed. (*C*) Representative nucleus of 3D multicolor DNA FISH using probes for the transfected BAC backbone (red) and *FBXO32* region (*FBXO32*, light blue) in myoblasts transfected with BAC 4q-D4Z4n. Nuclei are counterstained with DAPI (blue). All images at 63X magnification. Scale bar=5 µm. n, number of nuclei analyzed. (*D*) Bar plots showing expression levels of *FBXO32, TRIB3* and *ZNF555* (FSHD1 lost-activated genes) in CN (grey) and FSHD1 (blue) myoblasts transfected with Ctrl BAC and BAC 4q-D4Z4n. Data were normalized on *GAPDH* expression. n=at least 3 (with the exception of *TRIB3* and *ZNF555* CN Ctrl BAC, n=2). S.e.m. is indicated. *P* values were calculated by paired one-tailed *t*-test with confidence interval of 99%. Dots represent the values of each replicate; asterisks represent statistical *P* values; for *FBXO32* Ctrl BAC vs. BAC 4q-D4Z4n in CN *p*=0.0182; for *FBXO32* Ctrl BAC vs BAC 4q-D4Z4n in FSHD1 *p*=0.0073; for *TRIB3* Ctrl BAC vs BAC 4q-D4Z4n in FSHD1 *p*=0.0281. (*E*) Bar plot showing expression levels of *LZTS3* (FSHD1 lost-repressed gene) in CN (grey) and FSHD1 (blue) myoblasts transfected with Ctrl BAC and BAC 4q-D4Z4n. Data were normalized on *GAPDH* expression. n=3 (with the exception of CN Ctrl BAC, n=2). S.e.m. is indicated. *P* value was calculated by paired one-tailed *t*-test with confidence interval of 99%. Dots represent the values of each replicate; asterisks represent statistical *P* values; for Ctrl BAC vs. BAC 4q-D4Z4n in FSHD1 *p*=0.0296. (*F*) Bar plots showing expression levels of *FOXO3* and *MYOG* (not interacting genes) in CN (grey) and FSHD1 (blue) myoblasts transfected with Ctrl BAC and BAC 4q-D4Z4n. Data were normalized on *GAPDH* expression. n=at least 3. S.e.m. is indicated. Dots represent the values of each replicate.

Collectively, these results demonstrate that the 4q-D4Z4 array directly modulates the transcription of its interacting targets, suggesting a simultaneous fine-tuning of genes that occupy the same topological domain.

## Discussion

Here, we sought to identify the mechanisms by which the contraction of the tandem repeat D4Z4 on Chromosome 4 contributes to FSHD pathogenesis, using an integrated multi-omics approach (4C-seq, ChIP-seq and RNA-seq).

We found that 4q-D4Z4 interactome is altered in FSHD1 patients. In particular, normal 4q-D4Z4 array contacts several regions in a peripheral nuclear domain, controlling their transcription (Fig. 6A). In FSHD1 patients, the shortened and hypomethylated 4q-D4Z4 array causes an impairment of the chromatin conformation, which results in the loss of contacts with atrophic genes, with their consequent chromatin structure alteration and transcriptional upregulation (Fig. 6B). In this regard, it is already demonstrated that chromatin topological structures predominantly consist of simultaneous multiplex chromatin interactions with high heterogeneity between individual cells (Jiang et al. 2016; Zheng et al. 2019). Indeed, we show that an ectopic wild type 4q-D4Z4 array has the ability to get in close spatial proximity to the endogenous locus, resulting in the restoration of the expression of multiple targets, opening the possibility for further mechanistic studies on the dynamics of 3D interactions.

**Figure 6.**
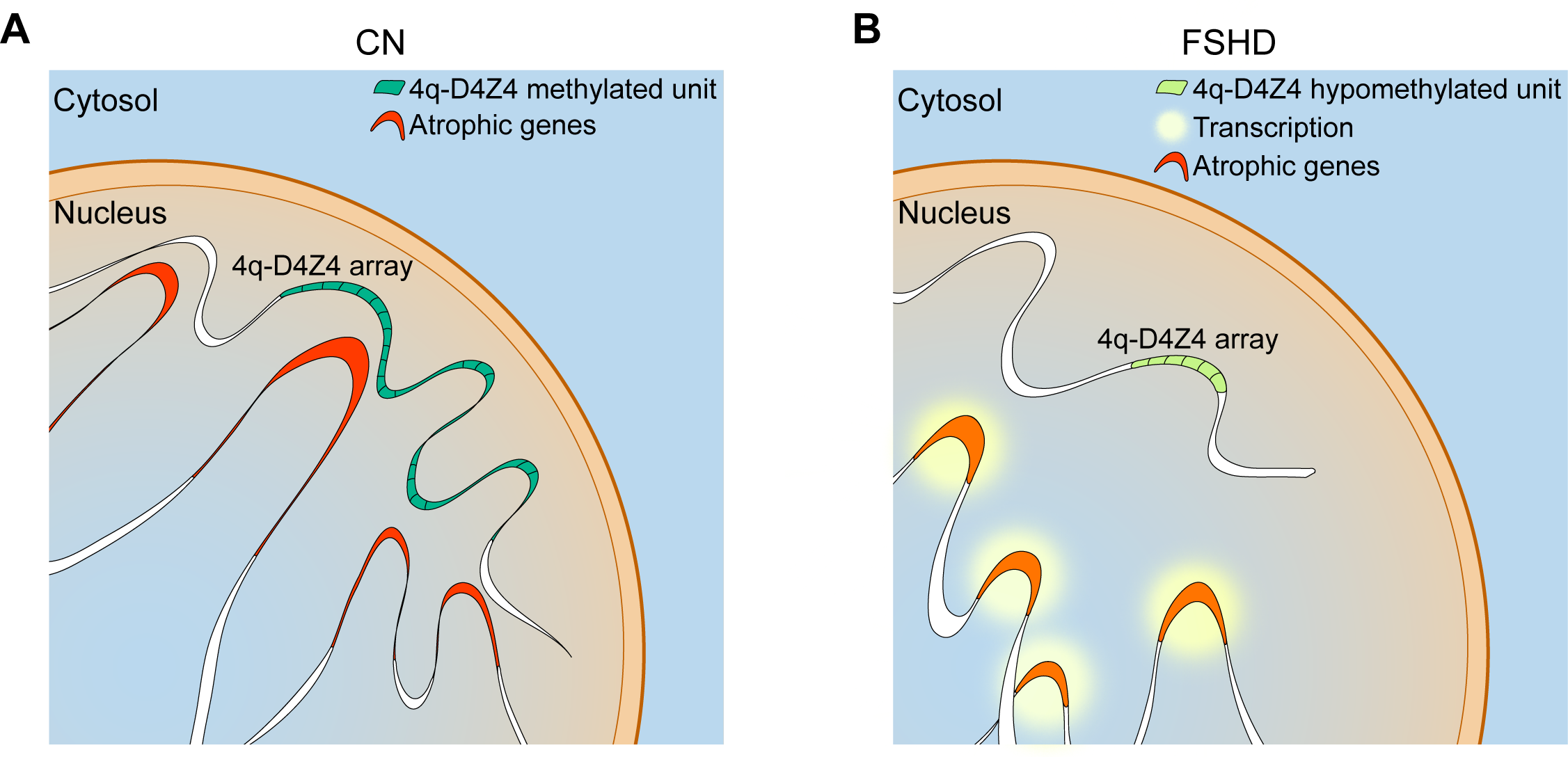
Model of 4q-D4Z4 mediated regulation of atrophic genes transcription. (*A*) 4q-D4Z4 array is interacting with a subset of atrophic genes, organizing their chromatin structure and keeping on hold their transcription in healthy donor muscle cells. (*B*) In FSHD1 patients’ muscle cells, the deleted and hypomethylated 4q-D4Z4 array causes an impairment of D4Z4 interactome leading to a chromatin switch towards an active state (mainly enhancer and promoter regions), which in turn results in the transcriptional upregulation of the atrophic genes.

We propose that the genetic deletion of 4q-D4Z4 array in FSHD1 patients leads to a rewired interactome that may represent an additional component of FSHD pathophysiology.

Since we discovered that the subset of genes losing contact with the 4q-D4Z4 array in FSHD1 mainly show chromatin state switches towards activation, we hypothesize that this might be consistent with a broader derepression occurring at the 4q-D4Z4 array, such as lesser PRC1/2 recruitment together or not with an enhanced activity of Trithorax complex, as already demonstrated *in cis* (Cabianca et al. 2012). Of note, SMCHD1 protein, mutated in FSHD2 patients (Lemmers et al. 2012), is now better characterized and involved in higher order chromatin organization of the inactive X Chromosome (Jansz et al. 2018; Wang et al. 2018). We could hypothesize that this architectural protein could have a central role in regulating 4q-D4Z4 interactions and that its altered function in FSHD1 (due to the contraction and hypomethylation of the array) and FSHD2 patients (due to its mutation) could explain the common atrophic signature. Indeed, FSHD-associated atrophy is one of the main signs of the disease (Tawil and Van Der Maarel 2006; Lanzuolo 2012; Sacconi et al. 2015), for which a direct link with the genetic defect remained elusive till now.

Importantly, we identified a core set of impaired atrophic genes, which is aberrantly transcribed in FSHD1 muscle cells used in this study and in other FSHD1 cells (Yao et al. 2014). Furthermore, they are also deregulated in FSHD2 muscle cells, indicating that the atrophic signature is a common trait in FSHD pathology. We further investigated *FBXO32* gene regulation, which was one of the top ranking; although it was already described overexpressed in muscle biopsies derived from FSHD1 fetuses and adults (Broucqsault et al. 2013), here we linked its transcriptional deregulation to the reduction in *FBXO32*/4q-D4Z4 interaction. This was predominantly observed in FSHD myoblasts compared to myotubes, in line with previous reports that changes in 3D structure precedes changes in gene expression (Hug et al. 2017; Krijger and de Laat 2017; Cheutin and Cavalli 2018) and already demonstrated also for *FRG1* gene (Bodega et al., 2009).

DNA repetitive elements are involved in a plethora of regulatory mechanisms, such as nuclear structure organization and spatiotemporal gene expression regulation (Gregory 2005; de Laat and Duboule 2013; Bodega and Orlando 2014). Additionally, recent studies have highlighted the contribution of satellite repeats in shaping 3D-genome folding and function, as evidenced for pericentromeric satellites (Politz et al. 2013; Wijchers et al. 2015) and DXZ4 macrosatellite (Giacalone et al. 1992; Rao et al. 2014; Deng et al. 2015; Darrow et al. 2016; Giorgetti et al. 2016). Our study is the first demonstration of a role of DNA repetitive elements in the alteration of genomic architecture in the context of a human genetic disease. It further corroborates the concept that perturbations of the 3D-genome structure are involved in various diseases (Krijger and de Laat 2016; Lupianez et al. 2016), such as cancers (Corces and Corces 2016; Rivera-Reyes et al. 2016; Achinger-Kawecka and Clark 2017) and developmental defects (Woltering et al. 2014; Lupianez et al. 2015; Woltering and Duboule 2015).

Our work highlights a novel role of DNA repeats in orchestrating gene transcription by shaping 3D genomic and chromatin architecture. We propose that perturbation of this DNA repeat-mediated regulatory network may be important in other complex genetic and epigenetic diseases.

## Methods

### Cell cultures

Although it was not always possible to ascertain the status of the muscle origin used in this study (see Supplemental Table S1, sheet “Cell line information”), the majority of the cells used derived from quadriceps, which in general was asymptomatic. Human primary myoblast cell lines from healthy donors (CN), patients affected by FSHD1 or FSHD2 were obtained from the Telethon BioBank of the C. Besta Neurological Institute, Milan, Italy and the Fields Center for FSHD of the Rochester Medical Center Dept. of Neurology, New York, USA; whereas human immortalized myoblast cell lines from healthy donors and FSHD1 patients were obtained from the University of Massachusetts Medical School Wellstone center for FSH Muscular Dystrophy Research, Wellstone Program & Dept. of Cell & Developmental Biology, Worcester, MA USA. Details of all cell lines are reported in Supplemental Table S1; details on media preparation and FACS analysis for Desmin staining are provided in Supplemental Methods.

### 4C-seq assay

The 4C assay was performed as previously described (Splinter et al. 2012) with minor modifications. A paired-end 4q-D4Z4-specific 4C-sequencing strategy was developed, where one 4C primer was designed to read the single sequence length polymorphism (SSLP) sequences located shortly upstream (almost 3.5 Kb) of the first D4Z4 repeat on 4q or 10q-D4Z4 arrays (Lemmers et al. 2007) and the second 4C primer reads into the captured sequence ligated to the ‘bait’ fragment. (Supplemental Fig. S1; Supplemental Table S1). Two donor muscle cell lines of CN (CN-3, CN-4) and FSHD1 (FSHD1-3, FSHD1-4) human primary myoblasts (3.5 × 10^6^ per sample) nuclei were processed. Five biological replicates (start to finish experiments) for each cell line were performed (Supplemental Fig. 3A). For *FBXO32* 4C-seq, we designed specific 4C primers as indicated in Supplemental Table S1. Two donor muscle cell lines of CN (CN-3, CN-4) and FSHD1 (FSHD1-3, FSHD1-4) human primary myoblasts (3.5 × 10^6^ per sample) nuclei were processed. From one to two biological replicates (start to finish experiments) for each cell line were performed. Hind III and Dpn II were used for enzymatic digestions. 4C samples were amplified using the bait and the SSLP specific primers. 4C sequencing libraries were prepared with 4C templates using the NEBNext Ultra DNA Library Prep Kit for Illumina, according to the manufacturer’s protocol, cleaned with Agencourt AMPure XP PCR Purification and sequenced on the Illumina NextSeq 500. For more details, see Supplemental Methods.

### 4C-Seq analysis

The paired-end 4q-D4Z4-specific 4C-seq reads were de-multiplexed based on the 4C bait reading primer that included the restriction site sequence. All reads were then trimmed and read pairs belonging to 4q-D4Z4, 10q-D4Z4 and 4q alleles were identified using SSLP reading mate (Read 1) (Supplemental Fig. S1; Supplemental Table S1) where no mismatch for the genotype sequence was allowed. Read pairs reading into the captured sequences ligated to the bait (Read 2) from the biological replicates of each donor muscle cell line were then pooled and mapped with Bowtie 2 (Langmead and Salzberg 2012). To find chromosome-wide interacting domains, 4C-ker (Raviram et al. 2016) was used. Reproducibility between donor muscle cell lines and quality of sequencing were assessed using Pearson correlation and cis/overall ratio (see Supplemental Methods). High frequency interactions for each viewpoint were intersected using BEDTools v2.2.4 (Quinlan and Hall 2010) and overlapping regions between donor muscle cell lines (CN-3 vs CN-4 and FSHD1-3 vs FSHD1-4) after removing overhangs were considered as high-confidence interacting domains. The interacting genes were defined as those that fall within the coordinates of these domains. Comparative analyses were performed between the 4q-D4Z4 alleles interactomes and also between the 4q and 10q-D4Z4 interactomes (see Supplemental Methods).

The paired-end *FBXO32* 4C-seq reads were demultiplexed based on the 4C bait reading primer that includes the restriction site sequence. All reads were trimmed and reads from the biological replicates of each donor muscle cell line were pooled and then mapped with Bowtie 2 (Langmead and Salzberg 2012). Reproducibility between donor muscle cell lines and quality of sequencing were assessed using Pearson correlation and cis/overall ratio. *Cis*-interacting domains were identified using 4C-ker (Raviram et al. 2016) and high-confidence interacting domains were selected. Full lists of interactions are available in Supplemental Table S2. For more details, see Supplemental Methods.

### ChIP-seq and ChIP-qPCR experiments

ChIP experiments were performed as previously described (Bodega et al. 2017) with minor modifications. The same donor muscle cell lines used for 4C-seq analysis of CN (CN-3, CN-4) and FSHD1 (FSHD1-3, FSHD1-4) human primary myoblasts and myotubes day 4 (3.5 × 10^6^ per sample) were processed for ChIP-seq analysis. For ChIP-seq and ChIP-qPCR, chromatin was immunoprecipitated with anti-H3K36me3 (ab9050, Abcam), anti-H3K4me1 (07- 436, Millipore), anti-H3K27ac (07-360, Millipore), anti-H3K4me3 (07-473, Millipore) and anti-H3K27me3 (07-449, Millipore); anti-RNA polymerase II CTD repeat YSPTSPS (phospho S5) antibody [4H8] (ab5408, Abcam). ChIP sequencing libraries were prepared using the NEBNext Ultra DNA Library Prep Kit for Illumina, according to the manufacturer’s protocol, cleaned with Agencourt AMPure XP PCR Purification and sequenced on the Illumina NextSeq 500 or Hiseq 2000. For ChIP-qPCR experiments, qRT-PCR analysis was performed on a StepOnePlus Real-Time PCR System, using power SYBR Green q-PCR master mix. The relative enrichment obtained by using all the antibodies was quantified after normalization for input chromatin. Primers used are reported in Supplemental Table S6. For more details, see Supplemental Methods.

### ChIP-seq analysis

We generated ChIP-seq datasets for CN myoblasts (MB) and myotubes (MT) day 4 for the following histone marks: H3K36me3, H3K4me3 and H3K27me3. H3K36me3, H3K4me1, H3K27ac, H3K4me3 and H3K27me3 datasets were generated for FSHD1 myoblasts and myotubes day 4. The following already published ChIP-seq datasets from ENCODE were used: H3K4me1 of human skeletal myoblasts (ENCSR000ANI), H3K27ac of human skeletal myoblasts (ENCSR000ANF), H3K4me1 of human skeletal myotubes (ENCSR000ANX) and H3K27ac of human skeletal myotubes (ENCSR000ANV). Reads were mapped with Bowtie 2 (Langmead and Salzberg 2012) on quality-checked (FastQC v0.11.2) and trimmed reads (trimmomatic v0.32; (Bolger et al. 2014)). For visualization of ChIP-seq tracks of independent samples, reads were normalized using bins per million mapped reads (BPM), same as TPM in RNA-seq, and to ensure fair comparison between all datasets, were further normalized to the respective input to produce coverage files reporting the log_2_ ratio of normalized read number between samples and inputs using bamCompare module. Quality and reproducibility assessment were done using deepTools2 package. For more details, see Supplemental Methods. Details on DUX4 ChIP-seq analysis is provided in Supplemental Methods.

### Chromatin state analysis

We used ChromHMM (Ernst and Kellis 2012) with default parameters to derive genome-wide chromatin states maps of CN and FSHD1 myoblasts and myotubes. We used the 5 histone marks H3K36me3, H3K4me1, H3K27ac, H3K4me3 and H3K27me3, as well as the respective input files, and binarized the data with BinarizeBed. We chose 15 states as the optimal number according to the maximal informative annotated genomic features and minimal redundancy. Subsequent functional annotations were attributed to each state choosing names and a color code for visualization according to the Roadmap Epigenomics Consortium nomenclature (Roadmap Epigenomics Consortium et al. 2015). Total number of derived chromatin features was similar between the samples (CN MB: 503,507; FSHD1 MB: 565,653; CN MT: 609,058; FSHD1 MT: 581,946). We performed overlap enrichment of the 15 chromatin states with known genome organization features (Supplementary Fig. S8A,B) and intersected chromatin states and genes bodies retrieved from GENCODE version 19 using BEDTools v2.2.4 (Quinlan and Hall 2010). Calculations of pairwise Jaccard were performed with BEDTools.

### Chromatin state switches analysis

In order to define whether CN and FSHD1 cells showed differences at gene chromatin state level, we took CN data as reference to search for specific switches in FSHD1. We intersected chromatin states retrieved from gene bodies in CN MB, CN MT, FSHD1 MB and FSHD1 MT. We postulated that a given state in a particular condition (CN MB or MT) should intersect another state in the other condition (FSHD1 MB or MT) in a reciprocal manner. We thus performed BEDTools intersect using –f .60 –r thus requiring that at least 60% of a state in CN MB or MT recovered a state in FSHD1 MB or MT in a reciprocal manner. In this way, identical states in the CN versus FSHD1 comparison (conserved states in the CN/FSHD1 comparison) as well as different states in the CN versus FSHD1 comparison (switching states in the CN/FSHD1 comparison) were retrieved. We focused on chromatin state switches between conditions (Supplemental Table S4) and added directionality to the chromatin state switches (that we chose to be active or repressive switches). We grouped the states into 3 main categories: promoters, enhancers and enhancer priming. The states involved in each group, as well as the definition of the directional switches they are involved in are summarized in Fig. 2B. For each gene, we also summarized all chromatin states expressed as a percentage of coverage across the gene body and defined the state with the highest coverage as being the “major state” for a given gene. We grouped those major states into the 2 categories of active and repressed as indicated in Fig. 2B. To obtain the genes showing directional switches, genes activated should display one of the following features: i) major state transition from repressed to active ii) major active state with at least one additional chromatin state switch towards activation. On the contrary, genes repressed should display either i) transition from an active to a repressed major state ii) major repressive state with at least one additional chromatin state switch towards repression as defined in Fig. 2B.

### RNA-seq assay and data analysis

RNA-seq studies were performed on the same donor muscle cell lines used for 4C-seq and ChIP-seq analyses of CN (CN-3, CN-4) and FSHD1 (FSHD1-3, FSHD1-4) human primary myoblasts and myotubes day 4. Briefly, total RNA was isolated using the miRNA Tissue kit on an automated Maxwell RSC extractor, following the manufacturer’s instructions. RNA integrity was assessed on TapeStation. Subsequently, RNA for each donor muscle cell line was used to generate single-end 75-bp sequencing libraries with the TruSeq Stranded mRNA Library Prep Kit, according to the manufacturer’s protocol. Sequencing was performed on a NextSeq500.

In addition, published RNA-seq dataset GSE56787 (Yao et al. 2014) consisting of human primary healthy, as well as FSHD myoblasts/myotubes obtained from the University of Rochester bio-repository (http://www.urmc.rochester.edu/fields-center) were also analyzed.

All fastq files were analyzed with FastQC v0.11.2. Adapters were removed and trimming was performed with Trimmomatic with standard parameters. Reads mapping to the reference genome GRCh37/hg19 was performed with STAR 2.3.0e (Dobin et al. 2013). The reference annotation used was GENCODEv19 and normalized FPKM (fragments per kilobase of transcript per million mapped reads) values were obtained with Cuffdiff (Trapnell et al. 2013). Normalized FPKM were log_2_ transformed and a value of 1 was added to all FPKM values to finally obtain log_2_ (1+ FPKM) values used in downstream analyses (Supplemental Table S4; Supplemental Table S5). For more details, see Supplemental Methods.

### Gene Ontology analysis

Gene Ontology analysis was performed on protein-coding genes and retrieved from different analysis with the Cytoscape v3.2.0 (Shannon et al. 2003) plug-in ClueGO v2.1.5 (Bindea et al. 2009). Statistically enriched Biological Processes (updated on 04/18/2016) were functionally grouped according to their k-score, and the most significant GO term of each group was used as summarizing GO term for the group. Full lists of GO terms and associated genes are available in Supplemental Table S5.

### Gene Set Enrichment Analysis (GSEA)

GSEA was performed as described in (Subramanian et al. 2005). The gene set was represented by the 319 lost-activated genes or by the 131 lost-repressed genes (genes from Fig. 3A; Supplemental Table S5). We tested if those genes were significantly enriched in a gene expression dataset associated with a skeletal muscle atrophic condition (disuse muscle atrophy, GSE21496; Supplemental Table S5; (Reich et al. 2010)). We also tested the association of our gene sets with a gene expression dataset from skeletal muscle hypertrophy (GSE12474; Supplemental Table S5; (Goto et al. 2011)). GSEA was performed on those datasets with the ranking metric Signal2noise with 1,000 phenotype permutations for statistical assessment of enrichment.

### Three-dimensional multicolor DNA FISH

To produce probes for 3D multicolor DNA FISH we used the following BAC DNA clones (BACPAC Resources Program, CHORI): CH16-77M12 (D4Z4, containing at least 15 units of D4Z4 repeat, B Bodega, unpublished and (Cabianca et al. 2012)), RP11-279K24 (4q), RP11-846C19 (C-), RP11-115K4 (C+), RP11-288G11 (10q26.3) and RP11-174I12 (*FBXO32*). Probes used for 3D multicolor DNA FISH in transfection experiments presented in Fig. 5A-C were produced from PCR designed on the pTARBAC6 backbone of the transfected CH16-291A23 BAC (BAC 4q-D4Z4n), on the pBACe3.6 backbone of the transfected RP11-2A16 BAC (Ctrl BAC), on a 35 Kb genomic region of 4q35.1 (4q) and on a 35 Kb genomic region of *FBXO32*. Primers used are reported in Supplemental Table S6. 1-3 μg of BAC DNA or pooled PCR products were labelled with bio-dUTP, dig-dUTP or cy3-dUTP through nick translation. The 3D multicolor DNA FISH assay was performed accordingly to (Cremer et al. 2008) with minor adaptations. One to three donor muscle cell lines of CN and FSHD human primary myoblasts or myotubes day 4 were processed for each experiment. An Eclipse Ti-E (Nikon Instruments) microscope was used to scan the nuclei, with an axial distance between 0.2-0.25 μm consecutive sections. In order to automatically analyze 3D multicolor DNA FISH in fluorescence cell image z-stacks, we developed a tool in MATLAB. The tool, that we named NuCLεD (Nuclear Contacts Locator in 3D), is capable to automatically detect and localize fluorescent 3D spots in cell image stacks. Measurements retrieved are shown in Supplemental Table S3. Details on 3D multicolor DNA FISH protocol and NuCLεD algorithm description are provided in Supplemental Methods.

### Chromatin conformation capture (3C)

The 3C assay was performed as previously described (Cortesi and Bodega 2016) with minor adaptations. Two donor muscle cell lines of CN (CN-3, CN-4) and FSHD1 (FSHD1-3, FSHD1-4) human primary myoblasts (3.5 × 10^6^ per sample) nuclei were processed. One to two biological replicates for each cell line was done. Digestion was performed using Hind III. A reference template was generated by digesting, mixing and ligating a BAC covering the genomic region of interest (*FBXO32* region, RP11-174I12). 3C templates and the reference template were used to perform PCR analysis with DreamTaq DNA Polymerase using primers designed around the Hind III restriction sites present at *FBXO32* region and, as bait primer, the same of *FBXO32*-4C (Supplemental Table S6) on a Veriti 96-Well Thermal Cycler. The PCR products were densitometrically quantified using the ImageJ software. Data are presented as the ratio of amplification obtained with 3C templates in respect to the reference template. For more details, see Supplemental Methods.

### BAC transfection

BAC transfections were performed accordingly to (Montigny et al. 2003) with minor adaptations. CN and FSHD1 human primary and immortalized myoblasts were plated. The following day, BAC DNA (RP11-2A16, as control BAC, representative of an unrelated and not interacting genomic region, Chr 17q21.33, and CH16-291A23, containing at least 15 units of D4Z4 repeat, B Bodega, unpublished and (Cabianca et al. 2012)) were diluted in Opti-MEM with the addition of P3000 Reagent. Lipofectamine 3000 Reagent were diluted in Opti-MEM. After 5 min BAC DNA (plus P3000) and Lipofectamine preparations were gently mixed and incubated for 20 min at room temperature. Transfection complexes were then added to the cells and incubated at 37 °C for 48 h. The primer pairs used for PCR or qRT-PCR amplifications are shown in Supplemental Table S6. For more details on transfection efficiency and DNA extraction, see Supplemental Methods.

### Statistics and Bioinformatics

To determine the significance between two groups, we used Wilcoxon matched-pairs signed rank test, Student’s *t*-test or Fisher’s exact test, as reported in Figure legends; exact *P* values and exact types of tests used are specified in Figure legends. For correlation analysis, we used Pearson correlation; the exact values are specified in the figures. Multiple comparisons were done by two-way ANOVA followed by Bonferroni post-test correction; exact *P* values and types of tests used are specified in Figure legends.

For all statistical tests, the 0.05 level of confidence was accepted for statistical significance; statistical significance is denoted by asterisks in figures, where * represent p-value <0.05, ** represents <0.01, *** represents <0.001 and **** represents <0.0001.

All reads were assessed for quality using FastQC and processed using Trimmomatic. They were aligned to the human genome (hg19) using either Bowtie 2 (Langmead and Salzberg 2012) or STAR 2.3.0e (Dobin et al. 2013). Aligning to GRCh38 is expected to provide similar results, as only a small number of bases change genome-wide with the major difference between the releases is in centromere assembly (Guo et al. 2017), which is not the focus of our study.

### Data access

Circular chromosome conformation capture and sequencing data (4C-seq), chromatin immunoprecipitation and RNA sequencing data (ChIP-seq and RNA-seq) for the human samples have been submitted to the NCBI Sequence Read Archive (SRA, http://www.ncbi.nlm.nih.gov/sra/) under the accession number SRP117155.

## Acknowledgments

We thank C. Lanzuolo, D. Gabellini, V. Ranzani, M. R. Matarazzo and E. Battaglioli for their stimulating discussions and constructive criticisms of this manuscript. We gratefully acknowledge M. Mora from the Telethon BioBank of the C. Besta Neurological Institute, R. Tawil from the Fields Center for FSHD of the Rochester Medical Center Dept. of Neurology and J. Chen from the University of Massachusetts Medical School Wellstone center for FSH Muscular Dystrophy Research, Wellstone Program & Dept. of Cell & Developmental Biology for providing the human myoblasts. The authors acknowledge the scientific and technical assistance of the INGM Imaging Facility (Istituto Nazionale di Genetica Molecolare “Romeo ed Enrica Invernizzi” (INGM), Milan, Italy), in particular C. Cordiglieri, for assistance during 3D multicolor DNA FISH images acquisition. pCMV-Mock and pCMV-DUX4 plasmids was a kind gift of D. Gabellini. This work has been supported by the following grants to B.B.: EPIGEN Italian flagship program, Association Française contre les Myopathies (AFM-Telethon, grant nr 18754) and Giovani Ricercatori, Italian Ministry of Health (GR-2011-02349383).

## Author contributions

A.C. and M.P. and S.S, designed and performed experiments, analyzed the data and wrote the manuscript. F.M and E.S. performed experiments. F.G., L.A., G.O. developed the new algorithm for 3D multicolor DNA FISH image analysis. C.C. and G.S. performed RNA-seq library preparation, NGS data processing and sequencing. B.B. conceived this study, designed experiments, analyzed the data and wrote the manuscript.

## Disclosure declaration

The authors declare no competing financial interests.

## Notes

http://www.ncbi.nlm.nih.gov/sra/

