## Supplementary Information for "4q-D4Z4 chromatin architecture regulates the transcription of muscle atrophic genes in FSHD"

Alice Cortesi^1♯^, Matthieu Pesant^1♯^, Shruti Sinha^1♯^, Federica Marasca^1^, Eleonora Sala^1^, Francesco Gregoretti^2^, Laura Antonelli^2^, Gennaro Oliva^2^, Chiara Chiereghin^3,4^, Giulia Soldà^3,4^, and Beatrice Bodega^1*^.

(1)  Istituto Nazionale di Genetica Molecolare "Romeo ed Enrica Invernizzi" (INGM), Milan, Italy

(2)  CNR Institute for High Performance Computing and Networking (ICAR), Naples, Italy

(3)  Department of Biomedical Sciences, Humanitas University, Pieve Emanuele, Milan, Italy

(4)  Humanitas Clinical and Research Center, Rozzano, Milan, Italy

(♯) These authors contributed equally to this work

**Table of contents**

**Supplemental Material**

**Supplemental Material for “4q-specific D4Z4 interactome is deregulated in FSHD1 patients”**

**Supplemental Material for “Genes that show impaired 4q-D4Z4 interactions and activated chromatin state are enriched for atrophic transcriptional signature”**

**Supplemental Material for “4q-D4Z4 lost interacting *FBXO32/ATROGIN1* gene is deregulated in FSHD patients at chromatin and transcriptional level”**

**Supplemental Material for “Ectopic 4q-D4Z4 array restores the expression of FSHD1 lost interacting genes”**

**Supplemental Methods**

**Supplemental Figures**

**Supplemental Figure S1.** 4q-D4Z4-4C strategy

**Supplemental Figure S2.** Characterization of human primary control and FSHD1 muscle cells used for 4C, ChIP and RNA-seq

**Supplemental Figure S3.** 4q and 10q-D4Z4 4C-seq quality controls

**Supplemental Figure S4.** 4q and 10q-D4Z4 4C-seq interactions quality controls

**Supplemental Figure S5.** 3D multicolor DNA FISH additional controls

**Supplemental Figure S6.** 4C-seq analysis for 4q alleles and 10q-D4Z4 interactomes

**Supplemental Figure S7.** ChIP-seq quality controls

**Supplemental Figure S8.** Chromatin states analysis

**Supplemental Figure S9.** Additional information on FSHD1 altered genes

**Supplemental Figure S10.** Examples of chromatin state definition of known DUX4 targets and their transcriptional levels

**Supplemental Figure S11.** Additional data on 10q-D4Z4 FSHD1 lost genes and GSEA

**Supplemental Figure S12.** Additional information on *FBXO32*/4q-D4Z4 *trans* interaction

**Supplemental Figure S13.** Additional information on *FBXO32* 4C-seq quality controls and *FBXO32* chromatin states

**Supplemental Figure S14.** Additional controls for *FBXO32* gene regulation and BAC transfection

**Supplemental Tables are given as a separate Excel files due to their length**

**Supplemental Table S1.** Cell lines, SSLP and 4C-seq primers

**Supplemental Table S2.** 4C-seq interactions analysis

**Supplemental Table S3.** 3D multicolor DNA FISH measurements

**Supplemental Table S4.** Chromatin state switches and RNA-seq expression levels

**Supplemental Table S5.** Gene Ontology, GSEA and atrophic genes expression levels

**Supplemental Table S6.** Primers

**References for Supplemental Material**

**Supplemental Material**

**Supplemental Material for “4q-specific D4Z4 interactome is deregulated in FSHD1 patients”**

With our 4C-seq approach, we retrieved 244 and 258 interacting regions for CN and FSHD1 respectively, which is in line with reported number of interactions retrieved using 4C-ker ((Qiu et al. 2019) around 400 interactions; (Schmiedel et al. 2016) around 200 interactions). In addition, we identified 4q allele specific interactome (4qA and 4qB) in CN and FSHD1. 4qA allele interacted with 411 and 392 regions in CN and FSHD1, whereas 4qB with 425 and 369 in CN and FSHD1 (Supplemental Table S2). The resolution of trans interacting domains is around 1Mb (Supplemental Table S2), similar to that of HiC.

Landscape of interactions of the two 4q alleles is highly shared both for CN and for FSHD1 Supplemental Fig. S6A; Supplemental Table S2), in agreement with what has been recently published in cell fate commitment (Rivera-Mulia et al. 2018). In line with this observation, the two alleles displayed a 70% of similarity of lost interactions, while showing almost 30% of specificity (Supplemental Fig. S6B, Supplemental Table S2). However, given that the number of reads used for calling 4q allele specific interactions was essentially the half of those used to call 4q interactions, the resolution of this analysis is lower, and therefore, further experiments will be required to investigate the biological relevance of the specific versus shared 4q allele interactions in the context of FSHD1 pathogenesis. In agreement with this observation, we did not distinguish between the 4q alleles for downstream analyses.

To assess the specificity of the results obtained for the 4q-D4Z4 interactome, we also analyzed the 10q-D4Z4 interactome, finding that the 4q and 10q shared only a subset of their interactions both in CN and FSHD1 (Supplemental Fig. S6C; Supplemental Table S2). 10q interactome was also impaired in FSHD1 condition (Supplemental Table S2); however, the intersection between 4q and 10q lost interactions in FSHD1 showed a very poor overlap (Supplemental Fig. S6D; Supplemental Table S2), confirming the specificity of the 4q-D4Z4 lost interactions in FSHD1. It has already been reported that 4q and 10q subtelomeres occupy the same peripheral nuclear domain (Masny et al. 2004; Tam et al. 2004); indeed, we retrieved their interaction in 4C-seq, that was further validated in 3D multicolor DNA FISH both in CN and FSHD1 (Supplemental Fig. S6E-H), suggesting that this spatial proximity could be the reason why the 10q interactome was also impaired in FSHD1. Moreover, the observation that the 10q-D4Z4 array contacted different genomic loci is further contributing to the concept that 10q deletions are not associated with FSHD disease (Bakker et al. 1995; Lemmers et al. 2001; Zhang et al. 2001).

**Supplemental Material for “Genes that show impaired 4q-D4Z4 interactions and activated chromatin state are enriched for atrophic transcriptional signature”.**

In order to highlight whether FSHD1 condition could exert a specific chromatin state feature in respect to CN, we assessed the global extent of conservation and variability of the chromatin states between CN and FSHD1. For a given state we computed the Jaccard value of overlap of each CN versus FSHD1 pairwise comparison (i.e. 1_TssA from CN MB versus 1_TssA from FSHD1 MB, etc). Chromatin segmentation analysis revealed chromatin state transitions in FSHD1 cells at enhancers and promoters; in particular we found that promoter-flanking regions (states 2-4 and 12) and enhancers in general (states 7-11) displayed the lowest Jaccard values, indicative of variability for those states between CN and FSHD1 (Supplemental Fig. S8C).

Interestingly, we observed that 40% of FSHD1 lost-activated and 62% of FSHD1 lost-repressed genes maintained the directionality in their chromatin switches during differentiation (MB versus MT; Supplemental Fig. S9C); the percentage of lost genes showing opposite chromatin switches during differentiation was remarkably lower, ranging from 2% to 5% (Supplemental Fig. S9C).

FSHD1 transcriptional program is recognized to be highly contributed by the transcription factor DUX4 (Jagannathan et al. 2016). However, only 5% of FSHD1 altered genes (lost-activated and repressed) were regulated by the master regulator DUX4 and only 2% of DUX4 regulated genes (Jagannathan et al. 2016) are comprised within the FSHD1 altered genes (Supplemental Fig. S9D), suggesting that DUX4 may regulate a small subset of genes already altered in their chromatin structure during myogenic differentiation. ChIP-seq and ChromHMM tracks from CN and FSHD1 MB and MT illustrating chromatin states alterations for known DUX4 deregulated genes (Jagannathan et al. 2016) that are either activated or repressed by DUX4 are presented in Supplemental Fig. S10A-D and their corresponding transcriptional deregulation is showed in Supplemental Fig. S10E.

**Supplemental Material for “4q-D4Z4 lost interacting *FBXO32/ATROGIN1* gene is deregulated in FSHD patients at chromatin and transcriptional level“**

We further asked whether the master regulator DUX4 (Yao et al. 2014) was necessary and sufficient to modulate *FBXO32* expression. As DUX4 expression is increased during FSHD1 myogenic differentiation (Supplemental Fig. S10E; (Balog et al. 2015)), we overexpressed DUX4 in CN and FSHD1 myoblasts (Supplemental Fig. S14B,C). We did not detect any effect on *FBXO32* transcription, whereas the DUX4 target *RFPL2* was upregulated and conversely, *RPL13A* (not a DUX4 target gene) transcription was not affected (Supplemental Fig. S14D; (Ferri et al. 2015)).

**Supplemental Methods**

**Cell cultures**

Cell lines from the Telethon BioBank were cultured in Dulbecco's modified Eagle medium (DMEM) supplemented with 20% Fetal bovine serum (FBS), 2 mM L-glutamine, 10 μg/ml Human insulin, 25 ng/ml Human fibroblast growth factor (hFGF), 10 ng/ml Human epidermal growth factor (hEGF) (proliferating medium), and then induced to differentiate by means of DMEM supplemented with 5% Horse serum (HS) and 100 μg/ml Human insulin (differentiating medium). Cell lines from the Fields Center for FSHD were cultured in F10 Nutrient Mixture supplemented with 20% FBS, 2 mM L-glutamine, 10 ng/ml hFGF, 0.4 μg/ml Dexamethasone (proliferating medium), and then induced to differentiate by means of DMEM supplemented with 2% HS, 10 μg/ml Human insulin, 4.5 g/L D-Glucose and 0.11 g/L Sodium pyruvate (differentiating medium). Immortalized cell lines were grown in plates coated with 0.1% Gelatin and cultured in Media X (4:1 DMEM/Medium 199 plus 0.88 mg/L Sodium pyruvate and 3.4 g/L Sodium bicarbonate) supplemented with 15% FBS, 10 ng/ml hFGF, 2.5 ng/ml Human hepatocyte growth factor (hHGF), 0.055 μg/ml Dexamethasone, 0.03 μg/ml Zinc sulfate, 1.4 μg/ml Vitamin B12, 0.02 M HEPES (proliferating medium). All of the patients satisfied the accepted clinical criteria for FSHD. FSHD1 patients had undergone DNA diagnosis and were identified as carriers of small (<38 Kb, <11 repeats) 4q35-located D4Z4 repeat arrays, as determined by p13E-11 hybridization to Eco RI-digested and Eco RI/Bln I-digested genomic DNA. FSHD2 patients had undergone DNA diagnosis and were identified as carriers of normal (>38 Kb, >11 repeats) 4q35-located D4Z4 repeat arrays, as determined by p13E-11 hybridization to Eco RI-digested and Eco RI/Bln I-digested genomic DNA, with an hypomethylated status of the array. Details of all cell lines are reported in Supplemental Table S1.

**FACS analysis for Desmin staining**

CN and FSHD1 myoblasts were collected with trypsin and cross-linked in 1% formaldehyde (Sigma) in growth medium. Primary antibody staining was carried out overnight at 4 °C in MACS running buffer at dilutions of 1:100 (Desmin antibody (D33) MA5-13259). Cells were washed three times with MACS buffer, at 1,600 x g, 4 °C. Secondary antibody staining was carried out at 30 °C in MACS running buffer at a dilution of 1:200 for 30 min. Secondary antibodies were coupled to Alexa Fluor 488 (Thermo Fisher Scientific). Cells were washed three times with MACS buffer, 1,600 x g, 4 °C, suspended in 200 μL of PBS and analyzed at FACScanto (BD Biosciences). Mock samples have been treated with secondary antibody only.

**4C-seq assay**

The first enzymatic digestion was performed with 600 U of Hind III (NEB) overnight at 37 °C in agitation. DNA circles were purified by phenol-chloroform-alcohol isoamyl extraction and ethanol precipitation and then resuspended in 10 mM Tris-HCl pH 7.5. The second enzymatic digestion was performed with 50 U of Dpn II (NEB) overnight at 37 °C. Ligation was performed with 200 U of T4 DNA Ligase (NEB) in 14 ml 1× Ligation buffer overnight at 16 °C with gentle agitation. 4C samples were amplified in 25 μl PCR reactions (12.5 ng per reaction, 200 ng of 4C template total) using the bait and the SSLP specific primers (Supplemental Table S1) and Phusion High-Fidelity DNA Polymerase (Thermo Fisher Scientific). 4C sequencing libraries were prepared with 500 ng of 4C templates using the NEBNext Ultra DNA Library Prep Kit for Illumina (NEB) according to the manufacturer’s protocol, without size selection and 5 cycles of PCR. Finally, 4C sequencing libraries were cleaned with Agencourt AMPure XP PCR Purification (Beckman Coulter), eluted in TE (10 mM Tris-HCl pH 8 and 1 mM EDTA) and sequenced on the NextSeq 500 (Illumina).

**4C-Seq analysis**

The upstream region of the 4q-D4Z4 array is a complex and duplicated region, in particular, it presents more than 98% of identity with the proximal 40 Kb of 10q-D4Z4 array (Deidda et al. 1995). In order to discard the duplicated regions and to specifically distinguish 4q and 10q mediated interactions, we developed a paired-end 4q-D4Z4-specific 4C-sequencing strategy (see above). Taking advantage of the single sequence length polymorphism (SSLP) sequences located in the proximal 4q and 10q-D4Z4 arrays (Lemmers et al. 2007), we designed 4C-sequencing primers (Supplemental Fig. S1; Supplemental Table S1), such that one 4C primer reads in the SSLP sequences and the second 4C primer reads into the captured sequence ligated to the ‘bait’ fragment. The SSLP reading primer was used to categorize the read-pairs as 4q-D4Z4, 10q-D4Z4 and 4q alleles specific. The analysis was performed as detailed in the steps below.

1. All 4C-seq reads were de-multiplexed based on the 4C bait reading primer (Read 2) (Supplemental Table S1) that included the restriction site sequence where one mismatch was allowed for the reading primer and no mismatches were allowed for the restriction site sequence.
2. All reads were trimmed for sequencing primer excluding the restriction site sequence. Reads were then trimmed for low quality using Trimmomatic v0.32 (Bolger et al. 2014) and unpaired reads were removed.
3. 4C bait pairs belonging to 4q-D4Z4, 10q-D4Z4 and 4q alleles were identified using SSLP reading mate (Read 1) (Supplemental Fig. S1C, Supplemental Table S1) were no mismatch for the genotype sequence was allowed.
4. All reads from the biological replicates for each donor cell line were then pooled and mapped to a reduced genome derived from hg19 consisting of all unique 75 nt-long regions surrounding Hind III sites. In case of 4q alleles, reads from the biological replicates of all donor cell lines of CN (CN-3, CN-4) and FSHD1 (FSHD1-3, FSHD1-4) were then pooled to obtain optimal number of reads necessary for calling interactions with high confidence in CN and FSHD1 conditions and mapped to a reduced genome derived from hg19 consisting of all unique 75 nt-long regions surrounding Hind III sites. Reads were mapped with Bowtie 2 (Langmead and Salzberg 2012) and parameters -N = 0 and −5 6 was used to trim the read sequences.
5. Chromosome-wide interacting domains were identified in 4q-D4Z4, 10q-D4Z4 and 4q alleles 4C-seq using 4C-ker (Raviram et al. 2016). Parameters of k=12 for *cis*, k=4 for nearbait and k=19 for *trans* interactions were used for analyzing 4q-D4Z4, 10q-D4Z4 and 4q alleles, except for cis (k=10) in 10q-D4Z4 and nearbait (k=4) in 4q-D4Z4. Briefly, 4C-ker uses a 3 state HMM to define regions of high, low and no significant interactions. It uses the k-th nearest neighbor approach to determine the adaptive window sizes that accounts for the change in 4C-seq coverage in different genomic regions. The value of k determines the number of observed fragments to be analyzed within each window. Interactions identified as high frequency by 4C-ker were used for downstream analyses.
6. High-confidence interacting domains specific for CN and FSHD1 conditions were identified in 4q-D4Z4 and 10q-D4Z4 4C-seq. Briefly, the high frequency interactions called in the cell lines of the respective condition (CN-3, CN-4 for CN and FSHD1-3, FSHD1-4 for FSHD) were intersected using BEDTools v2.2.4 (Quinlan and Hall 2010) and the overlapping genomic regions in CN-3 vs CN-4 and FSHD1-3 vs FSHD1-4 after removing the overhangs were selected. The interacting genes were then defined as those that fall within the coordinates of these high-confidence interacting domains.

Quality assessment of 4C-seq was performed at different steps.

1. Assessment of erroneously distinguishing reads coming from 4q-D4Z4, 10q-D4Z4 and 4q alleles was performed using the formula below (Wang et al. 2012).

$${Error Rate}_{\mathrm{experiment}}= \frac{\sum E_{sslp}}{\sum R_{sslp}}$$

where, E_sslp_ is the number of reads identical to the generated erroneous SSLP sequences, containing minimum number of all possible errors (mismatch/dinucleotide insertion) needed for one sslp sequence to obtain another sslp sequence. and R_sslp_ denotes the true number of reads with the sslp sequences and is calculated as below.

$$R_{\mathrm{sslp}}= G_{sslp}+E_{sslp}$$

where G_sslp_ is the number of reads with the exact sslp sequence (with 0 mismatch) of the genotype of the sample (Supplementary Table S1). SSLP sequence are of nature (CA_n_AACA_n_/CA_n_), hence the minimum number of mutations needed for the wrong assignment of alleles are position specific. Examples: 4qA (161, CA_10_AACA_11_) can be wrongly identified as 4qB (163, CA_10_AACA_10_) if there is a mismatch at 43^rd^/44^th^ position; similarly, a minimum of one mismatch at the position 21/22 (NA/CN) followed by mismatch at the position 39/40 (NA/CN) will lead to wrongly distinguish allele 4qA (161, CA_10_AACA_11_) and 4qB (168, CA_19_); a minimum of one dinucleotide insertion before AA followed by a mismatch after 8 CA repeats: can lead to ineptly distinguish between 4qA (161, CA_10_AACA_11_) / 4qB (163, CA_10_AACA_10_) and 10q-D4Z4 (CA_9_AACA_8_) [error sequence [CA_n_](NN)AACA_8_[(C/N)(N/A)]CA_(1/2)_]; error in distinguishing 4qB (168, CA_19_) and 10q-D4Z4 (CA_9_AACA_8_) can happen due to a mismatch at position 19/20 (NA/CN) followed by mismatch at the position 35/36. On an average 0.6% error rate was observed in distinguishing between any two alleles (Supplemental Table S1).

To further assess inaccuracy in assigning sslp due to PCR or sequencing error rate, was calculated the error rate of assigning an sslp not expected to be present in a determined donor cell line.

$${Error Rate}_{\mathrm{sample}}= \frac{\sum E_{sslp}}{\sum R_{sslp}}$$

where, R_sslp_ denotes all the sslp sequences under study and E_sslp_ is the number of reads identical to the sslp not present in the sample. In principle, this leads to calculating error rate for CN-3, CN-4 and FSHD1-3 as the percentage of 168 sslp over all sslp reads (161,163,168 and 166) and for FSHD1-4 as the percentage of 163 sslp over all sslp reads (161,163,168 and 166). On an average 0.16% error rate was observed in wrongly identifying an sslp (Supplemental Table S1).

2. Reproducibility of 4C signals between the donor cell lines for each condition was assessed using Pearson correlation on running sum of reads for a window of 100 fragends (Supplemental Fig. S3B,D). Additionally one to one comparison among all donor cell lines was performed at the level of all fragends (Supplemental Fig. S3C,E).
3. Quality of the experiment was assessed using the cis/overall ratio criteria as proposed by (van de Werken et al. 2012) (Supplemental Fig. S3F,G).
4. In order to assess the overall effect of experimental covariates and batch effects on the identified interactions in the donor cell lines of 4q-D4Z4 and 10q-D4Z4 4C-seq, Principal Component Analysis (PCA) was performed (Supplemental Fig. S4A). Briefly, fragends present within interacting regions were assigned a value of one and others as zero. PCA was then performed on the fragends with the assigned values using prcomp in R package (R Core Team 2017).
5. To further ascertain the robustness of *trans* interactions of 4q-D4Z4 array in CN and FSHD1, hierarchical clustering was performed on normalized 4C-signal of *trans* interacting fragends (Supplemental Fig. S4B). Normalized 4C-signal was calculated as fragend per million mapped reads, a method similar to TPM in RNA-seq, on *trans* interacting fregends. Further, fragends with 1 standard deviation for the normalized 4C signal across the samples were used to perform the clustering.

For *FBXO32* 4C-seq, we designed specific 4C primers as indicated in Supplemental Table S1. Reads were processed and analyzed as described above for 4q-D4Z4 4C-seq. Reproducibility of 4C signals between the donor cell lines for each condition was assessed using Pearson correlation on running sum of reads for a window of 100 fragends (Supplemental Fig. S13B). Additionally, one to one comparison among all donor cell lines was performed at the level of all fragends (Supplemental Fig. S13C). Quality of the experiment was assessed using the cis/overall ratio criteria as proposed by (van de Werken et al. 2012) (Supplemental Fig. S13D). To find interacting domains near the *FBXO32* locus, 4C-ker (Raviram et al. 2016) was run using the parameter k=5 for nearbait. High-confidence interacting domains for CN and FSHD1 conditions were identified by intersecting high frequency interactions called in the cell lines of the respective condition (CN-3, CN-4 for CN and FSHD1-3, FSHD1-4 for FSHD1) using BEDTools v2.2.4 (Quinlan and Hall 2010) and the overlapping genomic regions in CN-3 vs CN-4 and FSHD1-3 vs FSHD1-4 after removing overhangs were selected.

For visualization of 4C signals, BedGraph tracks for normalized number of reads (reads per million, RPM) overlapping fragends were smoothed over three fragends by running mean. For regions close to the viewpoint, normalized number of reads (reads per million, RPM) overlapping fragends were smoothed over three fragends by running mean for the selected genomic region. For visualization of regions close to *FBXO32*, normalized bedgraph generated by 4C-ker was used. BedGraph files were visualized in the UCSC browser. Circos software (Krzywinski et al. 2009) was used to visualize genome-wide 4C interactions.

**4C-seq interactome comparison**

Interactomes were compared using intersectBed from BEDTools v2.2.4 (Quinlan and Hall 2010). Shared genomic intervals (exact coordinates) between the interacting regions after removing the overhangs were considered as common interactions. In order to identify the lost interactions in FSHD1 for the 4qA and 4qB alleles, first the interactions detected for CN from the two alleles were merged, using mergeBed from BEDTools v2.2.4 (Quinlan and Hall 2010). These interactions were then intersected with the interactions from FSHD1 4qA and 4qB alleles using intersectBed (parameter -v). Interactions not having any overlap with FSHD1 4qA and 4qB alleles interactions were considered as 4qA and 4qB lost interactions respectively. Lost interactions of 4qA and 4qB alleles were then compared to identify interactions lost from both the alleles and those specifically lost from each allele using intersectBed as described above.

**ChIP-seq and ChIP-qPCR experiments**

Cells were cross-linked in 1% formaldehyde and then lysed in 50 mM HEPES KOH, pH 7.5, 10 mM NaCl, 1 mM EDTA, 10% glycerol, 0.5% NP-40, 0.25% Triton X-100. Nuclei were pelleted, washed in 10 mM Tris-HCl, pH 8, 200 mM NaCl, 1 mM EDTA, 0.5 mM EGTA and lysed in 10 mM Tris-HCl, pH 8, 100 mM NaCl, 1 mM EDTA, 0.5 mM EGTA, 0.1% Na-deoxycholate, 0.5% N-laurylsarcosine. Chromatin was sheared (BRANSON A250) and immunoprecipitated. The immunocomplexes were recovered with magnetic Dynabeads (protein G; Invitrogen), washed and de-cross-linked overnight at 65 °C. ChIP sequencing libraries were prepared with 5 ng of ChIP templates using the NEBNext Ultra DNA Library Prep Kit for Illumina (NEB) according to the manufacturer’s protocol, without size selection and 12 cycles of PCR. Finally, ChIP sequencing libraries were cleaned with Agencourt AMPure XP PCR Purification (Beckman Coulter), eluted in 10 mM Tris-HCl pH 8 and sequenced on the NextSeq 500 or Hiseq 2000 (Illumina).

For ChIP qPCR experiments, qRT-PCR analysis was performed on a StepOnePlus Real-Time PCR System (Applied Biosystem), using power SYBR Green q-PCR master mix (Thermo Fisher Scientific).

**ChIP-seq analysis**

Reads mapping to the reference genome GRCh37/hg19 was performed with Bowtie 2 (Langmead and Salzberg 2012) on quality-checked (FastQC v0.11.2) and trimmed reads (trimmomatic v0.32 (Bolger et al. 2014)). To account for differences in sequencing depth between ENCODE datasets and ours, we subsampled H3K4me1 and H3K27ac FSHD1 ChIP-seq datasets to a total number of 20 million reads with samtools using the command samtools view –s 3.4 –b where 0.4 represents the fraction of subsampling for our samples. Mapped reads for the ChIP-seq datasets we generated ranged from 18-77 millions. For normalization and production of normalized signal tracks, we used the deepTools2 package (Ramirez et al. 2014).

Reproducibility between all datasets was assessed using the deepTools2 package. Pearson coefficient correlation was calculated and represented as heatmaps (Supplemental Fig. S7A) using the multiBamSummary with –binSize 10000 and plotCorrelation modules. PCA graph (Supplemental Fig. S7B) was obtained with the plotPCA module.

**ChIP-seq analysis of DUX4**

To access if FBXO32 is a target of DUX4, we used the published ChIP-seq dataset (Geng et al. 2012).  DUX4 (SRR372673, SRR372674 and SRR397888) and input (SRR372676) read files were downloaded from NCBI Sequence Read Archive (Kodama et al. 2012) in SRA file format. These were converted to fastq files using fastq-dump (NCBI SRA toolkit version 2.8.2).  All fastq files were checked for read quality with FastQC v0.11.2. Adapters were removed and trimming was performed with Trimmomatic (Bolger et al. 2014) with standard parameters as follows: ILLUMINACLIP (LEADING:3 TRAILING:3 SLIDINGWINDOWS:4:15 MINLEN:30). Reads were mapped to the reference genome GRCh37/hg19 using Bowtie 2 (Langmead and Salzberg 2012) with the preset parameters for --very-sensitive option. Peaks were called using MACS2 (Zhang et al. 2008) with default parameters and *P* value cutoff of 1e-6.

**RNA-seq data analysis**

Total RNA was isolated using the miRNA Tissue kit on an automated Maxwell RSC extractor (Promega), following the manufacturer's instructions. RNA integrity was assessed on a TapeStation (Agilent). Subsequently, 2 µg of RNA for each sample were used to generate single-end 75-bp sequencing libraries with the TruSeq Stranded mRNA Library Prep Kit (Illumina), according to the manufacturer's protocol. Sequencing was performed on a NextSeq500 (Illumina). Adapters were removed and trimming was performed with Trimmomatic with standard parameters as follows: ILLUMINACLIP (LEADING:3 TRAILING:3 SLIDINGWINDOWS:4:15 MINLEN:50). Reads mapping to the reference genome GRCh37/hg19 was performed with STAR 2.3.0e (Dobin et al. 2013). Normalized FPKM (fragments per kilobase of transcript per million mapped reads) values were obtained with Cuffdiff (Trapnell et al. 2013) using the parameters –library-norm-method geometric –emit-count-tables –multi- read-correct. Cuffdiff was run separately to analyze CN with FSHD1 and CN with FSHD2 in myoblasts, as well as myotubes. CN and FSHD1 are composed of all healthy and FSHD1 samples from both our datasets and Yao’s datasets (Yao et al. 2014) respectively.

**Total RNA extraction and cDNA synthesis**

Total RNA extraction was performed using the RNeasy Mini Kit (Qiagen) or the PureLink RNA Mini Kit (Thermo Fisher Scientific) according to the manufacturers’ protocols, and the purified RNA was treated with RNase-free DNase (Qiagen) or with PureLink DNase Set (Thermo Fisher Scientific) for 30 min to remove any residual DNA. RNA was reverse-transcribed using the SuperScript III First-Strand Synthesis SuperMix (Thermo Fisher Scientific) according to the manufacturer’s protocol. 1 ng of expected cDNA was used in qRT-PCR analysis for each reaction. qRT-PCR analysis was performed on an StepOnePlus Real-Time PCR System (Applied Biosystem), using power SYBR Green q-PCR master mix (Thermo Fisher Scientific). The relative expression of the investigated genes was obtained after normalization against glyceraldehyde 3-phosphate dehydrogenase (*GAPDH*) and then reported as absolute value or normalized value on myoblasts. The primer pairs used for real time amplifications are shown in Supplemental Table S6.

**Three-dimensional multicolor DNA FISH**

1-3 μg of BAC DNA or pooled PCR products were labelled with bio-dUTP (Thermo Fisher Scientific), dig-dUTP (Roche) or cy3-dUTP (Thermo Fisher Scientific) through nick translation in 50 μl of Labelling mix buffer (0.02 mM C-G-A dNTPs, 0.01 mM dTTP, 0.01 mM labelled dUTP, 50 mM Tris-HCl pH 7.8, 5 mM MgCl2, 10 mM b-mercaptoethanol, 10 ng/μl Bovine serum albumin (BSA), 0.05-0.1 U/μl DNA Polymerase I (Thermo Fisher Scientific), 0.004-0.001 U/μl Amplification Grade DNase I (Sigma)) for 30 min-2 h at 16 °C, to obtain an average probe size of 50 bp. Probes were collected by ethanol precipitation, resuspended in 10 mM Tris-HCl pH 7.5 and then quantified using a Nanodrop 1000 Spectrophotometer (Thermo Fisher Scientific). The 3D multicolor DNA FISH assay was performed accordingly to (Cremer et al. 2008) with minor adaptations. For a single experiment 100-300 ng of each probe was precipitated with 3.5 μg of Human Cot-1 DNA (Thermo Fisher Scientific) and 20 μg of Deoxyribonucleic acid, single stranded from salmon testes (Sigma), and then resuspended in 6 μl of Hybridization solution (50% formamide pH 7.0 (FA)/2× SSC/10% Dextran sulfate). Cells were plated directly on coverslips and fixed with 4% Paraformaldehyde (PFA) in 1× PBS and TWEEN 20 0.1% (PBS-T) for 10 min at room temperature at the state of myoblasts or after inducing the differentiation at day 4. During the last minute, few drops of 0.5% Triton X-100 in 1× PBS (PBS) were added and then cells were washed with 0.01% Triton X-100 in PBS three times for 3 min at room temperature. Cells were first permeabilized with 0.5% Triton X-100 in PBS for 10 min at room temperature. In order to remove RNA, samples were treated with RNase Cocktail Enzyme Mix (Thermo Fisher Scientific) for 1 h at 37 °C. Cells were subjected to other steps of permeabilization with 20% Glycerol in PBS overnight at room temperature, followed by four cycles of freeze and thaw interleaved by soak with 20% Glycerol in PBS. Permeabilized cells were washed with PBS three times for 10 min at room temperature. Cells were then incubated in 0.1 M HCl for 5 min at room temperature, followed by a rinse with 2× SSC and then incubated in 50% FA in 2× SSC for at least 30 min at room temperature. Slides were equilibrated in 2× SSC for 2 min, washed in PBS for 3 min and then treated with 0.0025-0.0075% pepsin in 0.01-0.03 N HCl for 2-4 min at room temperature to eliminate cytoskeleton. Pepsin was inactivated with 50 mM MgCl_2_ in PBS twice for 5 min. Nuclei were post-fixed with 1% PFA in PBS for 1 min, washed with PBS for 5 min and with 2× SSC twice, and then back to 50% FA in 2× SSC for at least 30 min at room temperature. Hybridization solution was loaded on a clean microscopic slide, coverslip with nuclei was turned upside down on the drop of hybridization mixture and sealed with rubber cement. Samples were denatured for 4 min at 75 °C and leaved to hybridize in a metallic box floating in a 37 °C water bath overnight. Samples were washed with 2× SSC three times for 5 min at 37 °C and with 0.1× SSC three times for 5 min at 60 °C, followed a rinse with 0.2% TWEEN 20 in 4× SSC. Aspecific binding sites were blocked with Blocking solution (4% BSA in 4× SSC, 0.2% TWEEN 20) for 20 min at 37 °C. Samples were then incubated in the appropriate concentration of Streptavidin, Alexa Fluor 647 conjugate (Thermo Fisher Scientific) (1:1,000) or DyLight 488 Labeled Anti-Digoxigenin/Digoxin (Vector Laboratories) (1:100) diluted in Blocking solution for 35 min in a dark and wet chamber at 37 °C. Samples were washed with 0.2% TWEEN 20 in 4× SSC three times for 3 min at 37 °C, equilibrated in PBS and post-fixed with 2% Formaldehyde in PBS for 10 min at room temperature. Finally, the 3D-fixed nuclei were washed with PBS three times for 5 min at room temperature, counterstained with 1 ng/μl DAPI in PBS for 10 min at room temperature and washed with PBS two times for 5 min at room temperature. Coverslips were mounted.

**NuCLεD (Nuclear Contacts Locator in 3D): the 3D multicolor DNA FISH analysis algorithm**

In pilot experiments (Supplemental Fig. S5A), 4q-D4Z4 3D multicolor DNA FISH were analyzed with the Volocity software, with the following pipeline used for each 3D-reconstructed field: Select nuclei in DAPI channel - ROI design around FISH specific signals of 4q-D4Z4 spots – Recognize objects in 488 channel as D4Z4 - Recognize objects in 568 channel as 4q - Set threshold for each channel - Measure distances from centroid of D4Z4 spots to centroid of 4q spots. In this way, for each identified nucleus, the distance between 4q-D4Z4 spots and 4q spots were calculated. For all the others 3D multicolor DNA FISH, we developed and used the 3D multicolor DNA FISH analysis tool NuCLεD.

In order to automatically analyze 3D multicolor DNA FISH in fluorescence cell image z-stacks we developed a tool in MATLAB. The tool, that we named NuCLεD (Nuclear Contacts Locator in 3D), is capable to automatically detect and localize fluorescent 3D spots in cell image stacks. To achieve this objective, the algorithm performs the 2D segmentation of cell nuclei and the detection of spots for each slice of the stack followed by the 3D reconstruction and identification of nuclei and spots. It then measures the relative positioning of spots in the nucleus and inter-spots distances, which greatly enrich our understanding of the 3D spatial organization of the spots within cell nucleus. The tool implements the following algorithm:

for each slice n of the stack

nuclei_n_ = **nuclei_seg**(I_dapi,n_) %Performs 2D nuclei segmentation

for each fluorescent field f=ATROGIN1,4q,10q

spot_f,n_ = **detect_spot**(I_f_) %Performs 2D spot detection

spot_vol_f_(:,:,n) = spot_f,n_(:,:)

endfor

nuclei_vol(:,:,n) = nuclei_n_(:,:)

endfor

nuclei_CC = **bwconncomp**(nuclei_vol)

nuclei_L = **labelmatrix**(nuclei_CC)

compute volume for each nucleus object in nuclei_CC

exclude nuclei whose volume is less than 10% of mean volumes

(Arredondo et al.)_M_ <- identified 3D nuclei

for each nucleus m in (Arredondo et al.)_M_

for each fluorescent field f=ATROGIN1,4q,10q

NCL_m_.spot_f_ <- detected spots within the 3D nucleus NCL_m_

spot_CC_f_ = **bwconncomp**(NCL_m_.spot_f_)

compute volume for each spot object in spot_CC_f_

exclude spots whose volume is below the SD of the volumes

{SPT_f_}_nf_ <- identified 3D spots

endfor

if number of identified spots nf = 2 for each f=ATROGIN1,4q,10q

nucleus m is deemed suitable for analysis

compute distances of any spot from any other spot

compute distances of any spot from the nuclear periphery

compute distances of any spot from the nuclear centroid

compute distance from nuclear periphery to nuclear centroid

endif

endfor

The function *nuclei_seg* performs a partition of cell image in nuclei regions and background implementing a region based segmentation algorithm (Goldstein T et al. 2010). The function *detect_spot* has four major steps. It first filters the image, applying the Laplacian of Gaussian (LoG) operator (*fspecial* MATLAB function) of size 13 and standard deviation 7; this enhances the signal in the areas where objects are present. Then the function applies the h-dome transformation (Vincent 1993) that extracts bright structures by cutting off the intensity of height h from the top, around local intensity maxima; we used h=0.5 with a neighborhood size of 19x19. We decided to not use a global operator after having observed that a spot in one part of the image could be lighter or darker than the background in another part. This is due to the facts that spots have inhomogeneous intensity distribution over the image and that the image may have an uneven background. In the third step, the function performs a thresholding on h-domes image that excludes pixels whose intensity values are below a threshold. The threshold is 1.96 standard deviations above the mean of domes intensity values. We therefore assumed that spot areas have significant intensity disparity with respect to other bright areas present in cell nucleus. Lastly, the function applies a thresholding operation based on the surface areas of the spots, in order to discard too small objects, which are probably just noise. It filters out spots smaller than a surface area of 25. *detect_spot* produces an accurate set of spots. *spot_volf* and *nuclei_vol* are 3D arrays that contain the positions of the detected spots and nuclei from all slices. 3D reconstructions of nuclei are obtained through the connected components algorithm (*bwconncomp* MATLAB function, using a connectivity of 26). 3D nuclei are then labelled by applying the *labelmatrix* MATLAB function so they can be easily separated each from the others. The tool computes volume of each 3D reconstruction, discarding objects whose volume is less than 10% of mean volumes which are just noise. 3D reconstructions of spots are obtained through the connected components algorithm (*bwconncomp* MATLAB function, using a connectivity of 26). Then a threshold operation is performed on the 3D spots to obtain the more significant ones: it keeps in all the spots whose volume is above the threshold of the standard deviation of the volumes. For each nucleus, the tool checks the number of identified spots for each fluorescent field. If there are exactly two spots for each fluorescent field, that nucleus is deemed suitable for the analysis: the tool computes the distances between the centroid of each spot, the distances of the centroid of each spot from the nuclear periphery and the nuclear centroid and the distance from the nuclear periphery to the nuclear centroid. Three-dimensional distances between specified genomic loci and the nuclear centroid were normalized on the maximum radius for each nucleus.

**Chromatin conformation capture (3C)**

Digestion was performed using 600 U of Hind III (NEB) at 37°C overnight with constant agitation. DNA fragments were purified by phenol-chloroform extraction and ethanol precipitation and then resuspended in 10 mM Tris-HCl pH 7.5. 3C templates and the reference template were used to perform PCR analysis on a Veriti 96-Well Thermal Cycler (Applied Biosystem), using DreamTaq DNA Polymerase (Thermo Fisher Scientific).

**Plasmid transfection**

For DUX4 transfection, CN and FSHD1 human primary and immortalized myoblasts were plated at 5 x 10^5^ cells/well in growth medium without antibiotics in 6-well dish containing a glass coverslip. Cells were leaved to adhere for 4-5 h prior transfection. 4 μg of plasmid DNA (pCMV-DUX4 or pCMV-Mock, kindly provided by Dr. Gabellini) were diluted in 250 μl final volume of room-temperature Opti-MEM (Thermo Fisher Scientific) and 10 μl Lipofectamine 2000 Reagent (Thermo Fisher Scientific) were diluted in 250 μl final volume of Opti-MEM. After 5 min plasmid DNA and Lipofectamine preparations were gently mixed and incubated for 20 min at room temperature. Transfection complexes were then added to the cells (replaced with 2 ml of fresh medium) and incubated at 37 °C in 5% CO_2_ for 24 h.

**Immunofluorescence**

For immunofluorescence staining of DUX4 in CN and FSHD1 human primary and immortalized myoblasts transfected with plasmids containing or not DUX4 ORF (pCMV-DUX4 or pCMV-Mock, kindly provided by Dr. Gabellini) were fixed with 4% PFA in PBS-T for 15 min at room temperature and then washed with PBS twice. Cells were permeabilized with 0.5% Triton X-100 in PBS for 10 min at room temperature with gentle agitation and then washed with PBS three times. Aspecific binding sites were blocked with Blocking solution (4% BSA in PBS-T) for 30 min at room temperature with gentle agitation. Samples were incubated with the primary rabbit-DUX4 antibody directed against the C- terminal region of DUX4 (E5-5, ab124699, 1:200, Abcam), diluted in 4% BSA in PBS-T for 3 h at room temperature with gentle agitation. Samples were washed in PBS-T three times and then incubated with Goat anti-Rabbit IgG (H+L) Cross-Adsorbed Secondary Antibody, Alexa Fluor 488 (A-11008, 1:1,000, Thermo Fisher Scientific) diluted in 4% BSA in PBS-T for 1 h at room temperature with gentle agitation in the dark. Samples were washed with PBS-T three times for 5 min and counterstained with 1 ng/μl DAPI in PBS for 10 min at room temperature, then washed with PBS-T two times for 5 min and with PBS two times for 5 min at room temperature. Coverslips were mounted. An Eclipse Ti-E (Nikon Instruments) microscope was used to scan the cells. DUX4 immunofluorescences were analyzed with the NIS software, with the following pipeline used for each field: Select nuclei in DAPI channel – Select DUX4 positive cells in 488 channel – Count the number of nuclei and the number of DUX4 positive cells. From these values, the percentage of DUX4 positive cells was calculated.

**BAC transfection**

In order to set up BAC transfection on human primary and immortalized myoblasts, we transfected human primary myoblasts with increasing amount (1 μg – 2 μg) of the 4q-D4Z4 BAC and a plasmid containing EGFP (pLEGFP) on 5 x 10^4^ cells/well plated in 12-well dish. We counted the percentage of EGFP positive cells, retrieving 23% of transfection for 1 μg and 45% of transfection for 2 μg (Supplemental Fig. S14F).We extracted DNA and performed a semi-quantitative PCR (25 cycles) both on pLEGFP and BAC backbone regions, retrieving comparable amplification efficiency (Supplemental Fig. S14G). We chose the condition of 2 μg for the following experiments, estimating that also BAC transfection efficiency was around 45%.

CN and FSHD1 human primary and immortalized myoblasts were plated at 6 x 10^4^ cells/well in growth medium without antibiotics in 12-well dish. The following day, 2 μg of BAC DNA (RP11-2A16, as control BAC, representative of an unrelated and not interacting genomic region Chr 17q21.33, and CH16-291A23, containing at least 15 units of D4Z4 repeat, B Bodega, unpublished and (Cabianca et al. 2012)) were diluted in 50 μl final volume of room-temperature Opti-MEM (Thermo Fisher Scientific) with the addition of 4 μl P3000 Reagent (Thermo Fisher Scientific). 4 μl Lipofectamine 3000 Reagent (Thermo Fisher Scientific) were diluted in 50 μl final volume of Opti-MEM. After 5 min BAC DNA (plus P3000) and Lipofectamine preparations were gently mixed and incubated for 20 min at room temperature. Transfection complexes were then added to the cells (replaced with 600 μl of fresh medium) and incubated at 37 °C in 5% CO_2_ for 24 h. Transfection medium was then replaced with fresh medium and the cells were maintained under standard conditions for other 24 h.

**DNA extraction**

Total DNA extraction was performed in 200 μl DNA Extraction Buffer (20 mM Tris-HCl pH 8, 5 mM EDTA and 150 mM NaCl) with the addition of 0.5% SDS and 80 μg of Proteinase K (Sigma) overnight at 37 °C with gentle agitation. The day after DNA was purified by phenol-chloroform and phenol-chloroform-alcohol isoamyl extraction and ethanol precipitation, and resuspended in 10 mM Tris-HCl pH 7.5. Approximately 16 ng of DNA sample was used in PCR analysis for each reaction. PCR analysis was performed on a Veriti 96-Well Thermal Cycler (Applied Biosystem), using DreamTaq DNA Polymerase (Thermo Fisher Scientific). The PCR products were densitometrically quantified using the ImageJ software. The relative transfection efficiency of the BACs used (RP11-2A16 and CH16-291A23), was calculated normalizing BACs specific PCR amplification on a genomic control region PCR amplification and then reported as normalized on CN myoblasts transfected with control BAC. The primer pairs used for PCR amplifications are shown in Supplemental Table S6.

**Supplemental Figures**

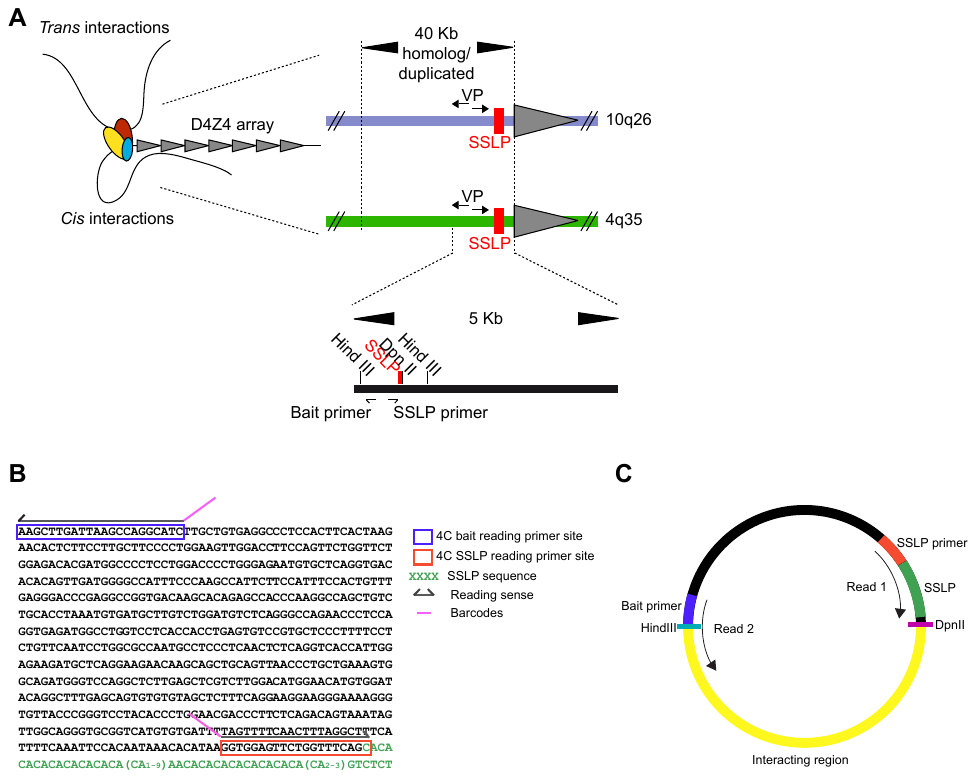

**Supplemental Figure S1. 4q-D4Z4-4C strategy**

(*A*) Scheme of the 4q-D4Z4 specific 4C-seq approach. The 40 Kb proximal region of the polymorphic 4q35.2 D4Z4 array displays more than 98% of sequence identity with 10q26.3. As a viewpoint (VP) a region within a window of 5 Kb upstream the first repeat of the 4q-D4Z4 array was used, nearby a single sequence length polymorphism (SSLP) that allowed to design a paired-end 4q-specific 4C (Bait primer and SSLP primer locations are represented). Paired-end reads discriminate the origin of interactions (Chr 4 and Chr 10) based on the SSLP sequence and retrieve the corresponding interacting region. (*B*) Sequence of the 4q-D4Z4 viewpoint region. 4C bait and SSLP reading primers sites are shown in blue and red boxes respectively; SSLP sequence is highlighted in green; arrows indicate the sense of the reading. (*C*) Representation of a 4C product. Read 1 refers to read pair 1, reading the SSLP sequence, and Read 2 refers to the corresponding mate, reading the interacting region.

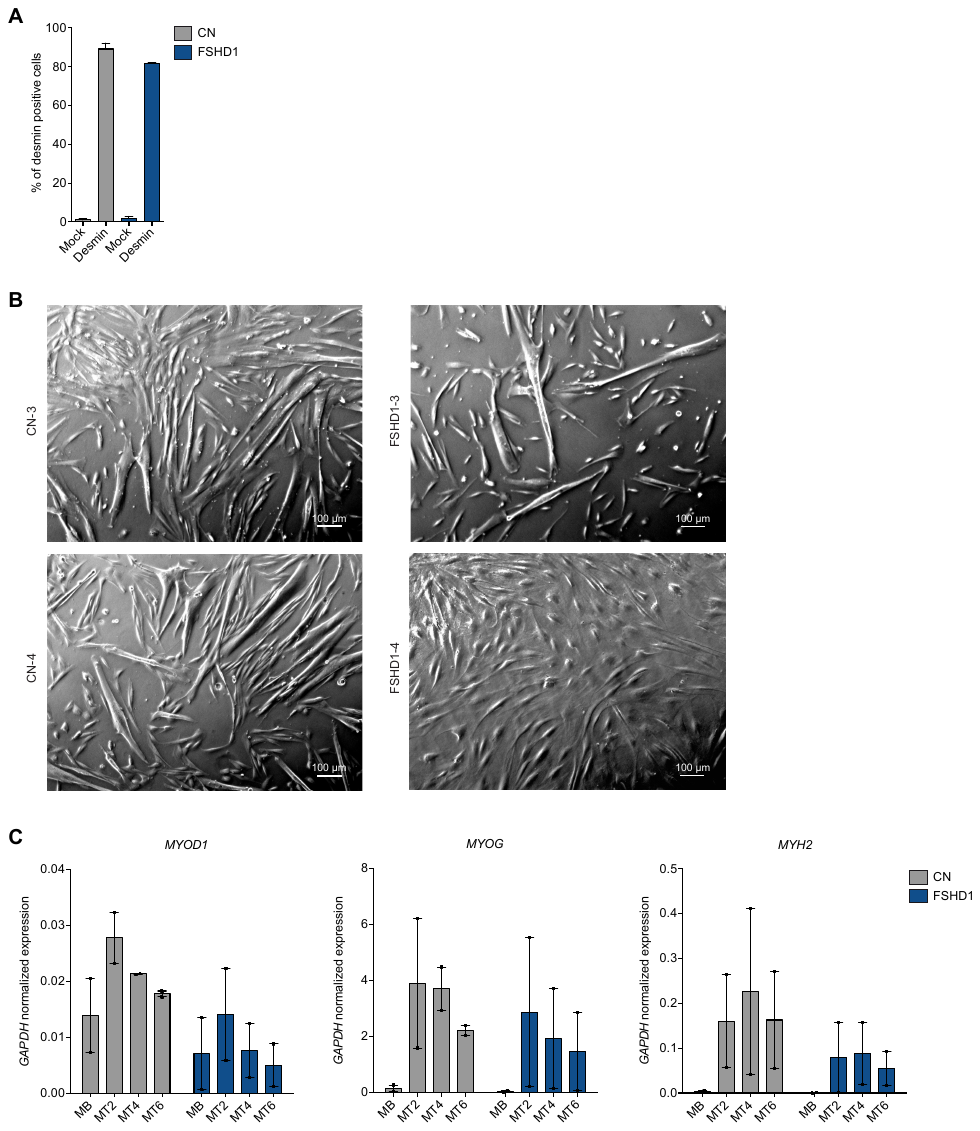

**Supplemental Figure S2. Characterization of human primary control and FSHD1 muscle cells used for 4C, ChIP and RNA-seq**

(*A*) Bar plot showing percentage of positive cells for desmin staining in CN (grey) and FSHD1 (blue) cells. 2 CN (CN-3, CN-4) and 2 FSHD1 (FSHD1-3, FSHD1-4). S.e.m. is indicated. (*B*) Light microscope images of myotubes day 4 (MT4) of CN-3, CN-4, FSHD1-3 and FSHD1-4 (*C*) Expression levels of *MYOD1, MYOG* and *MYH2* genes during CN (grey) and FSHD1 (blue) differentiation (MB, myoblasts, MT2, myotubes day 2, MT4, myotubes day 4, MT6, myotubes day 6). Data were normalized on *GAPDH* expression. n=2 CN (CN-3, CN4), FSHD1 (FSHD1-3, FSHD1-4). S.e.m. is indicated. Dots represent the values of each replicate.

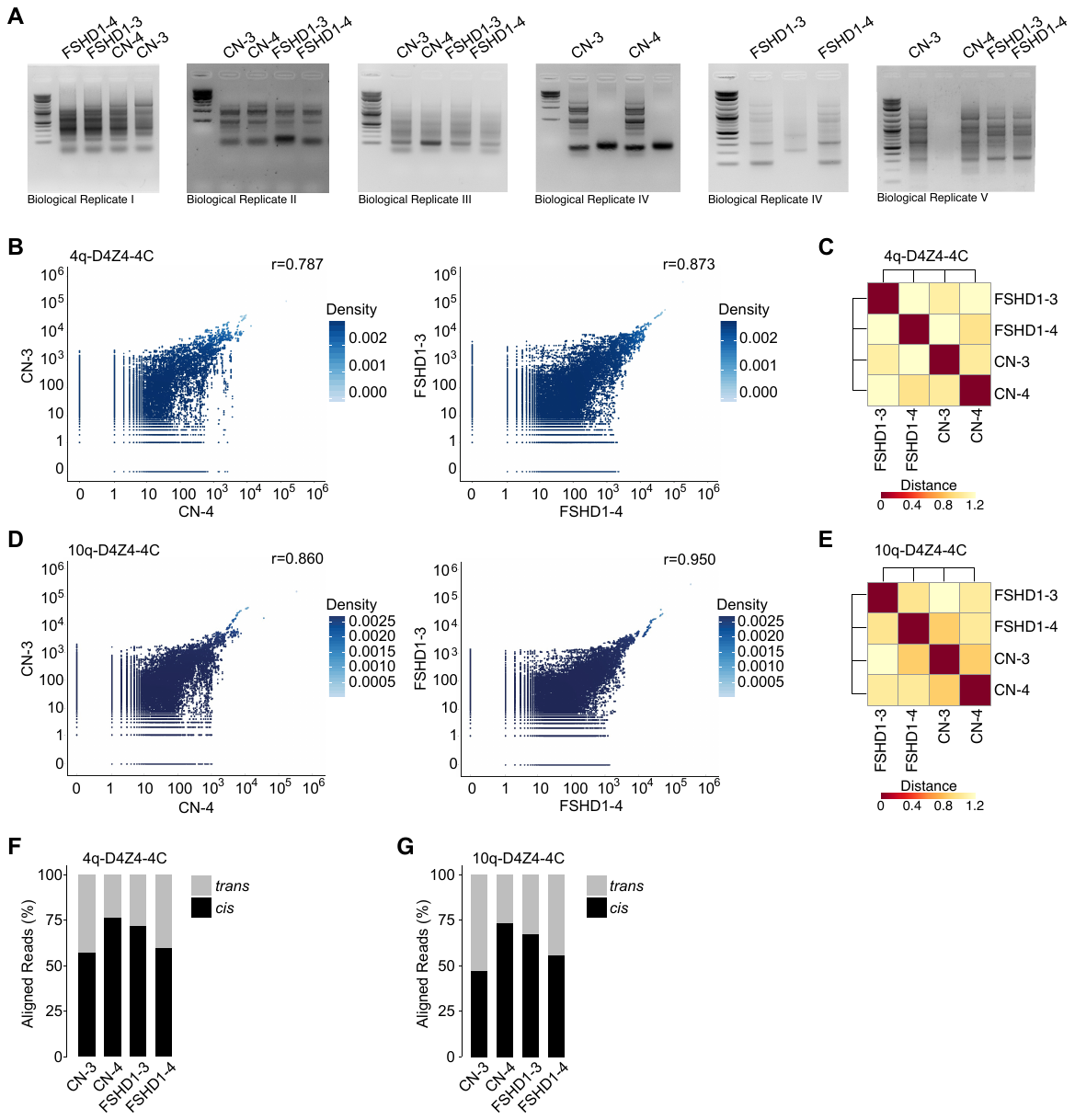

**Supplemental Figure S3. 4q and 10q-D4Z4 4C-seq quality controls**

(*A*) Agarose gel runs for 4q-D4Z4 4C PCRs for each replicate. (*B*) Scatter plots of running sum count reads for a window of 100 fragends for the 4q-D4Z4-4C. Pearson correlation coefficient shows the reproducibility between biological samples of CN (CN-3, CN-4) and FSHD1 (FSHD1-3, FSHD1-4). (*C*) Sample-to-sample comparison at the level of all fragends for the 4q-D4Z4-4C viewpoint. (*D*) Scatter plots of running sum count reads for a window of 100 fragends for the 10q-D4Z4-4C. Pearson correlation coefficient shows the reproducibility between biological samples of CN (CN-3, CN-4) and FSHD1 (FSHD1-3, FSHD1-4). (*E*) Sample-to-sample comparison at the level of all fragends for the 10q-D4Z4-4C viewpoint. (*F*) Bar plot showing percentages of mapped reads *in* *cis* and *in* *trans* for the 4q-D4Z4-4C viewpoint. (*G*) Bar plot showing percentages of mapped reads *in* *cis* and *in* *trans* for the 10q-D4Z4-4C viewpoint.

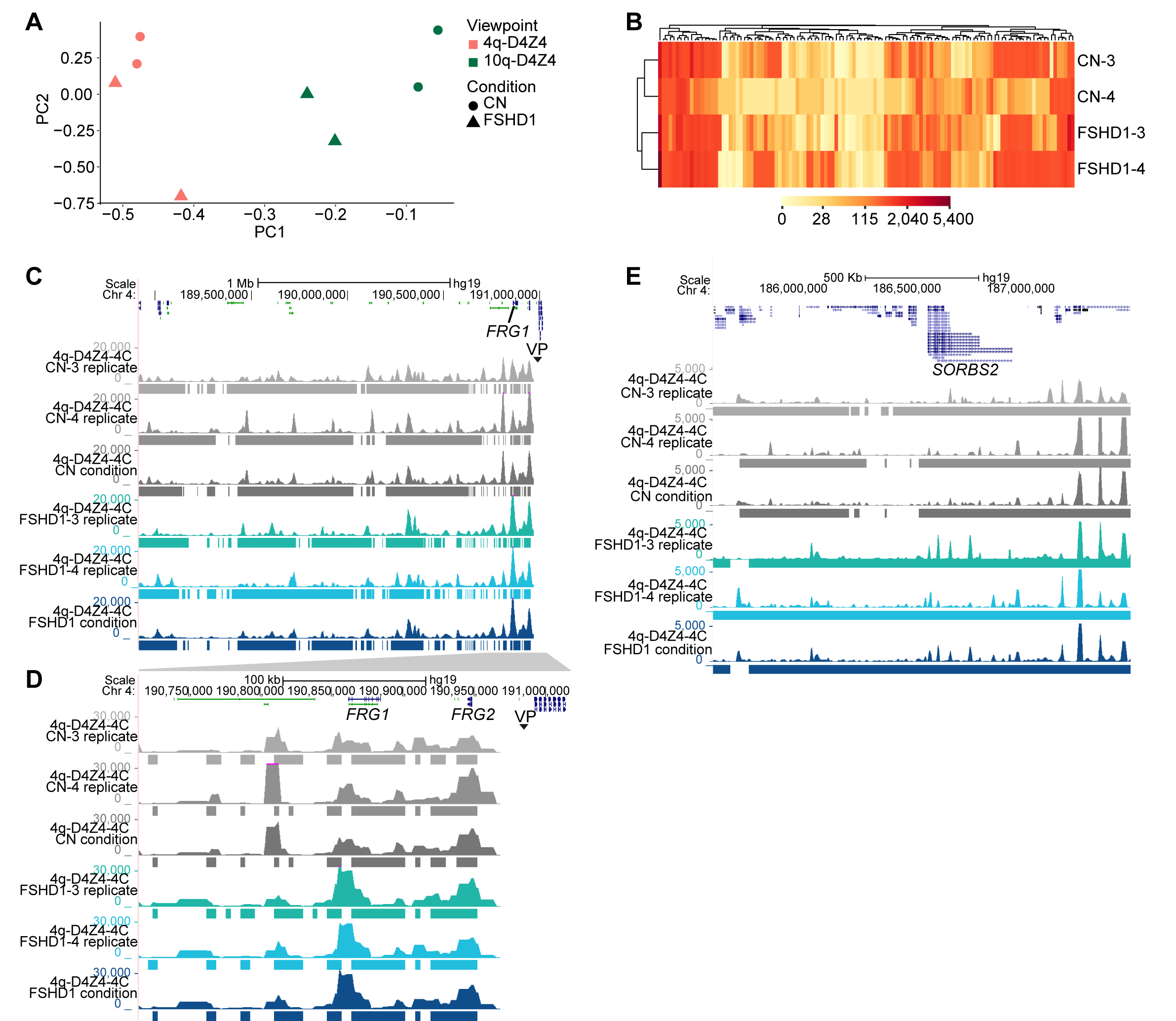

**Supplemental Figure S4. 4q and 10q-D4Z4 4C-seq interactions quality controls**

(*A*) Principal Component Analysis (PCA) representation of 4C-seq interactions for both the viewpoints (4q and 10q) in CN (CN-3, CN-4) and FSHD1 (FSHD1-3, FSHD1-4). (*B*) Hierarchical clustering of 4q-D4Z4 4C-seq signals for *trans* interacting fragends of CN (CN-3, CN-4) and FSHD1 (FSHD1-3, FSHD1-4). (*C*) 4q-D4Z4-4C normalized coverage tracks of a 2 Mb region upstream the 4q-D4Z4-4C VP in CN (grey) and FSHD1 (blue). Called 4C interacting regions for each donor as well as condition are presented. (*D*) Zoomed-in region of (*C*) showing 4q-D4Z4-4C normalized coverage tracks at the *FRG1* locus in CN (grey) and FSHD1 (blue). (*E*) Snapshot of *SORBS2* genomic region showing 4q-D4Z4-4C normalized coverage tracks at the locus in CN (grey) and FSHD1 (blue). Called 4C interacting regions for each donor as well as condition are presented under the respective 4C normalized coverage track.

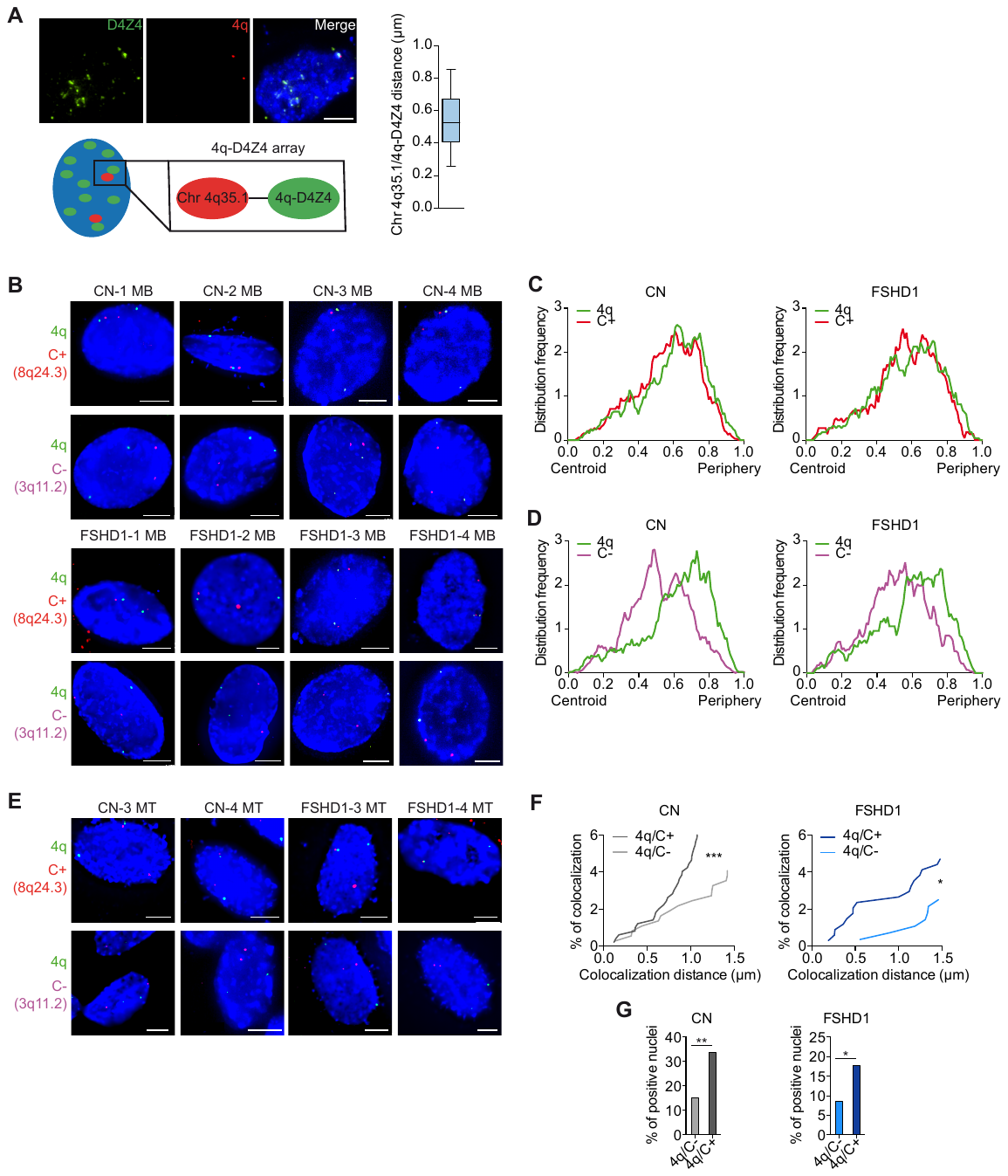

**Supplemental Figure S5. 3D multicolor DNA FISH additional controls**

(*A*) (left) Representative nucleus of 3D multicolor DNA FISH using probes mapping to D4Z4 repeat (green) and 4q35.1 region (4q, red) in CN cells and scheme of probes spots. Nuclei are counterstained with DAPI (blue). All images at 63X magnification. Scale bar=5 µm. (right) Box & whiskers plot showing the distribution of distances of 4q-D4Z4 spots from 4q35.1 region spots in CN and FSHD1 cells and whiskers extend to the 5-95 percentiles. (*B*) Representative nuclei of 3D multicolor DNA FISH using probes mapping to 4q35.1 region (4q, green), a 4q-D4Z4 positive interacting region (8q24.3, C+, red) and a 4q-D4Z4 not interacting region (3q11.2, C-, magenta) in CN (CN-1, CN-2, CN-3, CN-4) and FSHD1 (FSHD1-1, FSHD1-2, FSHD1-3, FSHD1-4) myoblasts. Nuclei are counterstained with DAPI (blue). All images at 63X magnification. Scale bar=5 µm. (*C*) Frequency distributions of normalized distances from the nuclear centroid of 4q35.1 region (4q, green) and positive interacting region (C+, red) in CN (left) and FSHD1 (right) myoblasts. n= 648 (CN) and 442 (FSHD1). (*D*) Frequency distributions of normalized distances from the nuclear centroid of 4q35.1 region (4q, green) and not interacting region (C-, magenta) in CN (left) and FSHD1 (right) myoblasts. n= 854 (CN) and 564 (FSHD1). (*E*) Representative nuclei of 3D multicolor DNA FISH using probes mapping to 4q35.1 region (4q, green), positive interacting region (8q24.3, C+, red) and a 4q-D4Z4 not interacting region (3q11.2, C-, magenta) in CN (CN-3, CN-4) and FSHD1 (FSHD1-3, FSHD1-4) myotubes. Nuclei are counterstained with DAPI (blue). All images at 63X magnification. Scale bar=5 µm. (*F*) Cumulative frequency distributions of distances (below 1.5 µm) between 4q and C+ and between 4q and C- (dark and light grey or blue respectively) in CN (left) and FSHD1 (right) myotubes. n= 496 (CN 4q/C+), 368 (CN 4q/C-), 340 (FSHD1 4q/C+) and 280 (FSHD1 4q/C-). *P* values were calculated by unpaired one-tailed *t*-test with confidence interval of 99%. Asterisks represent statistical *P* values; for 4q/C+ vs. 4q/C- in CN *p*=0.0001; for 4q/C+ vs. 4q/C- in FSHD1 *p*=0.0366. (*G*) Percentage of nuclei positive for the interactions (under the cut-off of 1.5 µm). n= 92 (CN 4q/C-), 124 (CN 4q/C+), 70 (FSHD1 4q/C-) and 85 (FSHD1 4q/C+). *P* values were calculated by fisher’s exact one-sided test with confidence interval of 99%. Asterisks represent statistical *P* values; for 4q/C- vs. 4q/C+ in CN *p*=0.0014; for 4q/C- vs. 4q/C+ in FSHD1 *p*=0.0483.

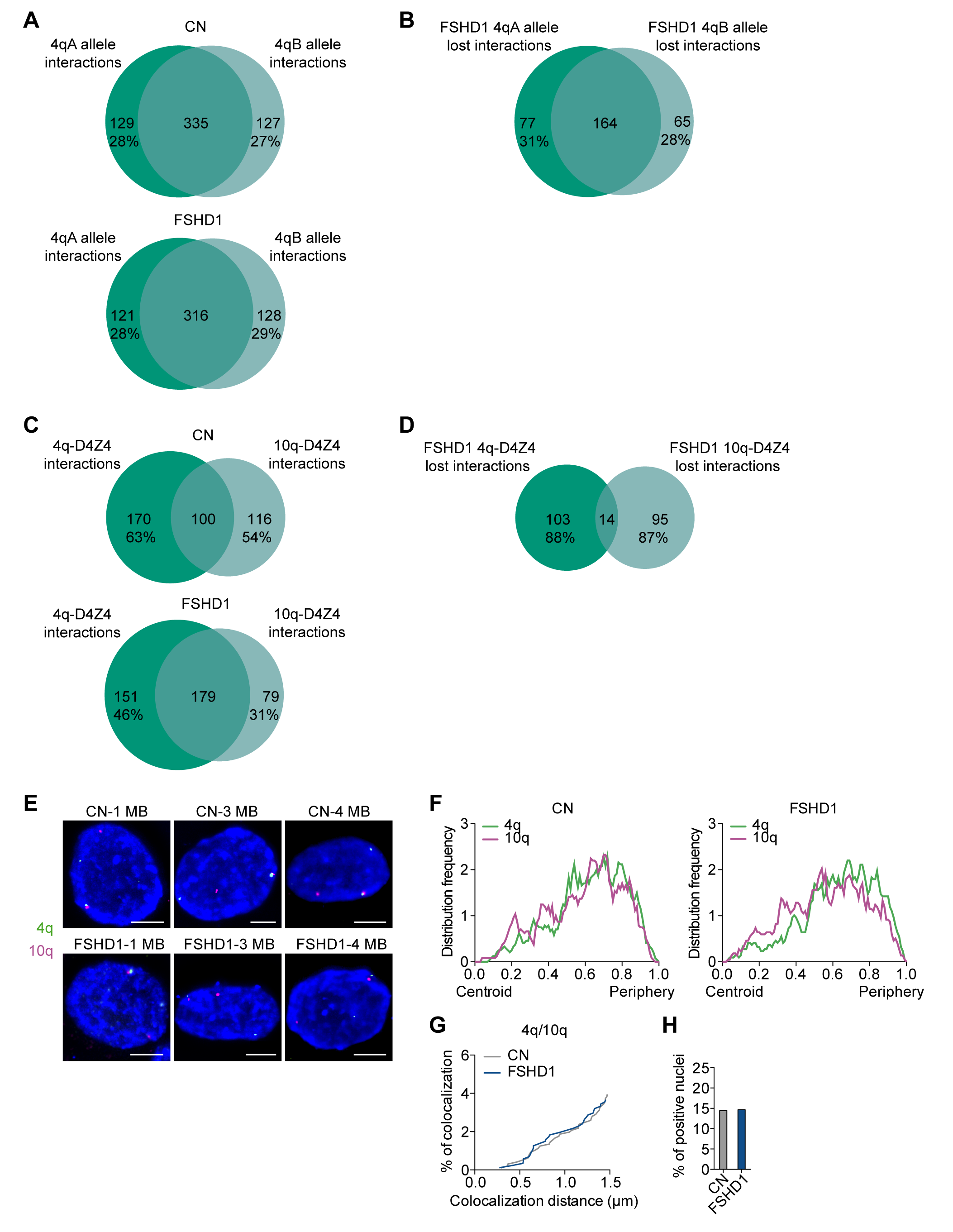

**Supplemental Figure S6. 4C-seq analysis for 4q alleles and 10q-D4Z4 interactomes**

(*A*) Venn representation of 4q-D4Z4 4qA and 4qB alleles interactomes, in CN (top) and FSHD1 (bottom). (*B*) Venn representation of 4q-D4Z4 4qA and 4qB alleles lost interactions comparison in FSHD1 cells. (*C*) Venn representation of 4q-D4Z4 and 10q-D4Z4 interactomes in CN (top) and FSHD1 (bottom). (*D*) Venn representation of 4q and 10q-D4Z4 lost interactions comparison in FSHD1 cells. (*E*) Representative nuclei of 3D multicolor DNA FISH using probes mapping to 4q35.1 region (4q, green) and 10q26.3 region (10q, magenta) in CN (CN-1, CN-3, CN-4) and FSHD1 (FSHD1-1, FSHD1-3, FSHD1-4) myoblasts. Nuclei are counterstained with DAPI (blue). All images at 63X magnification. Scale bar=5 µm. (*F*) Frequency distributions of normalized distances from the nuclear centroid of 4q35.1 region (4q, green) and 10q26.3 region (10q, magenta) in CN (left) and FSHD1 (right) myoblasts. n= 482 (CN) and 436 (FSHD1). (*G*) Cumulative frequency distributions of distances (below 1.5 µm) between 4q and 10q in CN (grey) and FSHD1 (blue) myoblasts. n= 964 (CN) and 872 (FSHD1). (*H*) Percentage of nuclei positive for the interactions (under the cut-off of 1.5 µm). n= 241 (CN) and 218 (FSHD1).

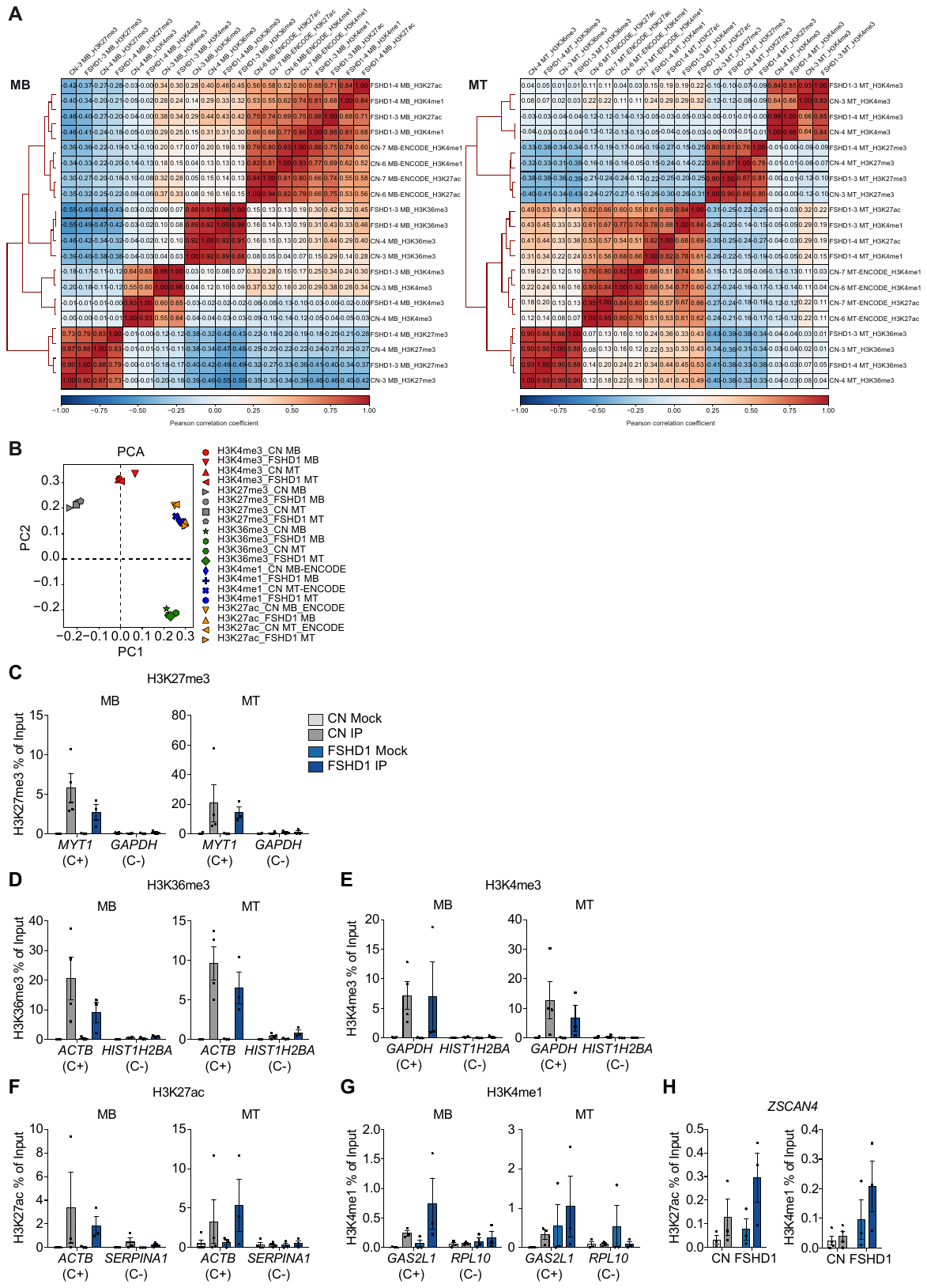

**Supplemental Figure S7. ChIP-seq quality controls**

(*A*) Clustered heatmaps of genome-wide enrichment signals of all the ChIP-seq datasets for each biological sample of CN and FSHD1 in MB (left) and MT (right). Samples were clustered according to Pearson correlation coefficient calculated in non-overlapping 10 Kb window. H3K4me1 and H3K27ac ChIP-seq datasets for CN MB/MT were from ENCODE (See Methods). (*B*) Principal Component Analysis (PCA) representation of genome-wide enrichment signals for all the histone marks we analyzed in CN and FSHD1 MB and MT. (*C*) Bar plots showing enrichment of H3K27me3 at positive (C+) and negative (C-) control region in MB (left) and MT (right) assessed by ChIP-qPCR experiment in CN (Mock in light grey and the IP in grey) and FSHD1 (Mock in light blue and the IP in blue) . Results are presented as % of Input. n=4 CN, 3 FSHD1. S.e.m. is indicated. Dots represent the values of each replicate. (*D*) Bar plots showing enrichment of H3K36me3 at positive (C+) and negative (C-) control region in MB (left) and MT (right) assessed by ChIP-qPCR experiment in CN (Mock in light grey and the IP in grey) and FSHD1 (Mock in light blue and the IP in blue) . Results are presented as % of Input. n=4 CN, 3 FSHD1. S.e.m. is indicated. Dots represent the values of each replicate. (*E*) Bar plots showing enrichment of H3K4me3 at positive (C+) and negative (C-) control region in MB (left) and MT (right) assessed by ChIP-qPCR experiment in CN (Mock in light grey and the IP in grey) and FSHD1 (Mock in light blue and the IP in blue). Results are presented as % of Input. n=4 CN, 3 FSHD1. S.e.m. is indicated. Dots represent the values of each replicate. (*F*) Bar plots showing enrichment of H3K27ac at positive (C+) and negative (C-) control region in MB (left) and MT (right) assessed by ChIP-qPCR experiment in CN (Mock in light grey and the IP in grey) and FSHD1 (Mock in light blue and the IP in blue). Results are presented as % of Input. n=3 CN, 3 FSHD1. S.e.m. is indicated. Dots represent the values of each replicate. (*G*) Bar plots showing enrichment of H3K4me1 at positive (C+) and negative (C-) control region in MB (left) and MT (right) assessed by ChIP-qPCR experiment in CN (Mock in light grey and the IP in grey) and FSHD1 (Mock in light blue and the IP in blue). Results are presented as % of Input. n=3 CN, 3 FSHD1. S.e.m. is indicated. Dots represent the values of each replicate. (*H*) Bar plots showing enrichment of H3K27ac (left) and H3K4me1 (right) at *ZSCAN4* region assessed by ChIP-qPCR experiment in CN (Mock in light grey and the IP in grey) and FSHD1 (Mock in light blue and the IP in blue) myoblasts. Results are presented as % of Input. n=3 CN H3K27ac, 3 FSHD1 H3K27ac, 4 CN H3K4me1 and 3 FSHD1 H3K4me1. S.e.m. is indicated. Dots represent the values of each replicate.

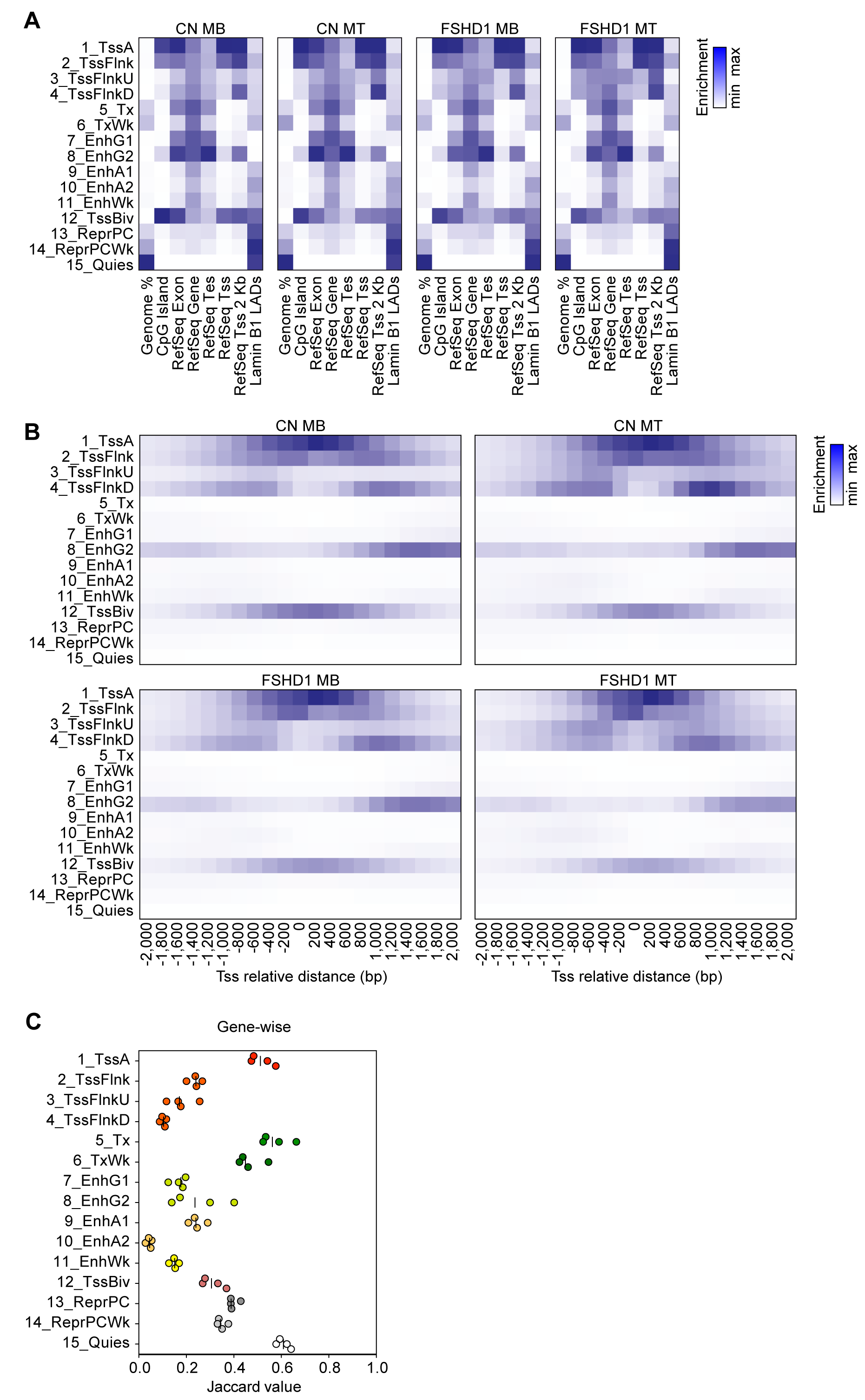

**Supplemental Figure S8. Chromatin states analysis**

(*A*) Heatmaps showing overlap of different genomic features (such as CpG islands, RefSeq Tss, RefSeq Tes and Lamin B1-Associated Domains) with chromatin states in CN MB/MT and FSHD1 MB/MT as determined with ChromHMM software. (*B*) Heatmaps showing enrichment of chromatin states around the Tss of RefSeq genes (+/- 2 Kb) in CN MB/MT and FSHD1 MB/MT. (*C*) Jaccard values of overlap of each CN versus FSHD1 pairwise comparison for a given state in both MB and MT at gene-wise level. Each dot shows one pairwise comparison (lines indicate medians).

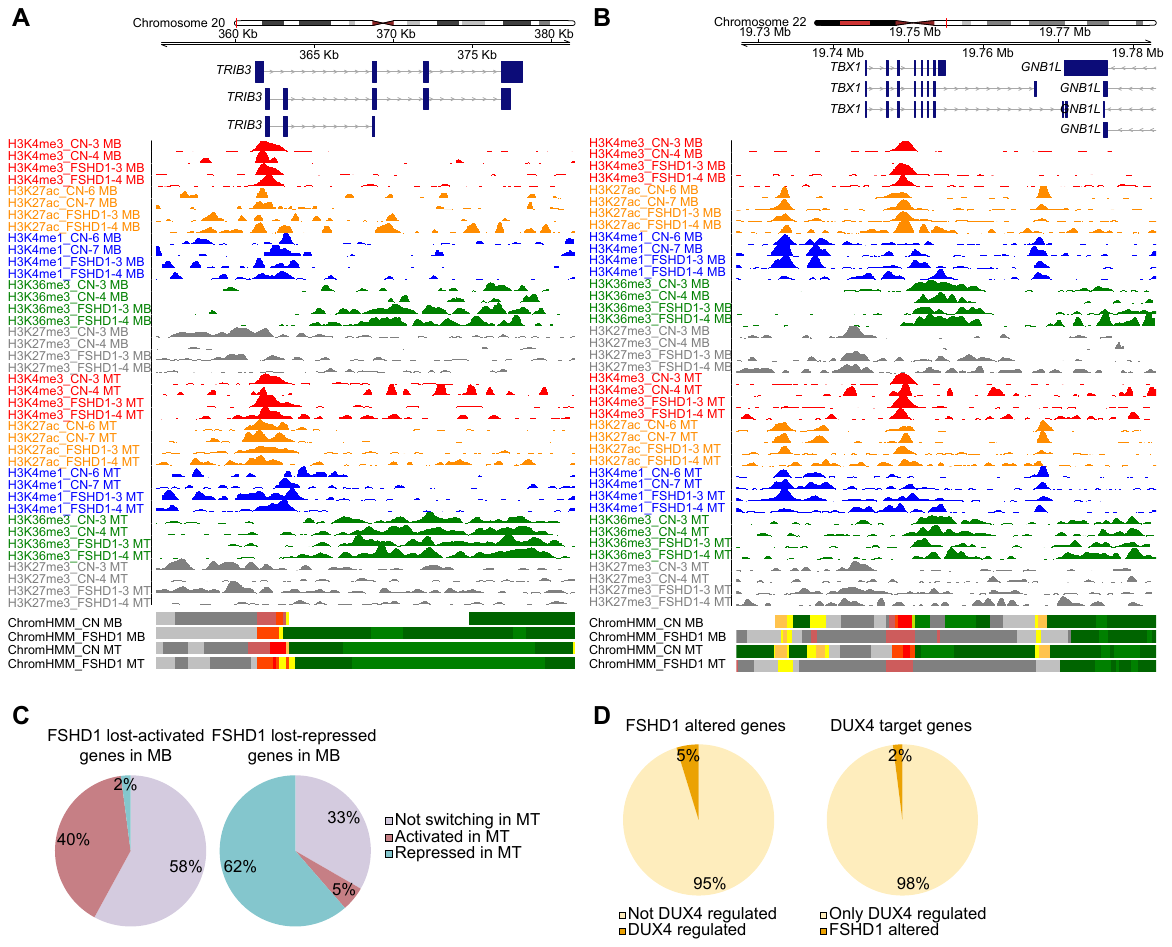

**Supplemental Figure S9. Additional information on FSHD1 altered genes**

(*A*) Representative ChIP-seq tracks of all histone modifications in each sample and chromatin states tracks of the FSHD1 lost-activated gene *TRIB3* of the GO term “proteasome-mediated ubiquitin-dependent protein catabolic process” from Fig. 3B. (*B*) Representative ChIP-seq tracks of all histone modifications in each sample and chromatin states tracks of the FSHD1 lost-repressed gene *TBX1* of the GO term “muscle tissue morphogenesis” from Fig. 3B. (*C*) Pie charts depicting percentages of altered genes in FSHD1 MB (activated on the left and repressed on the right) classified as not switching (violet), activated (red) and repressed (blue) in MT. (*D*) (left) Pie charts depicting percentages of altered genes in FSHD1 from Fig. 3A classified as DUX4-regulated (dark orange) or not (light orange) according to Jagannathan et al. (Jagannathan et al. 2016). (right) percentages of DUX4 target genes according to Jagannathan et al. (Jagannathan et al. 2016) classified as FSHD1 altered (orange) or only DUX4 regulated (yellow).

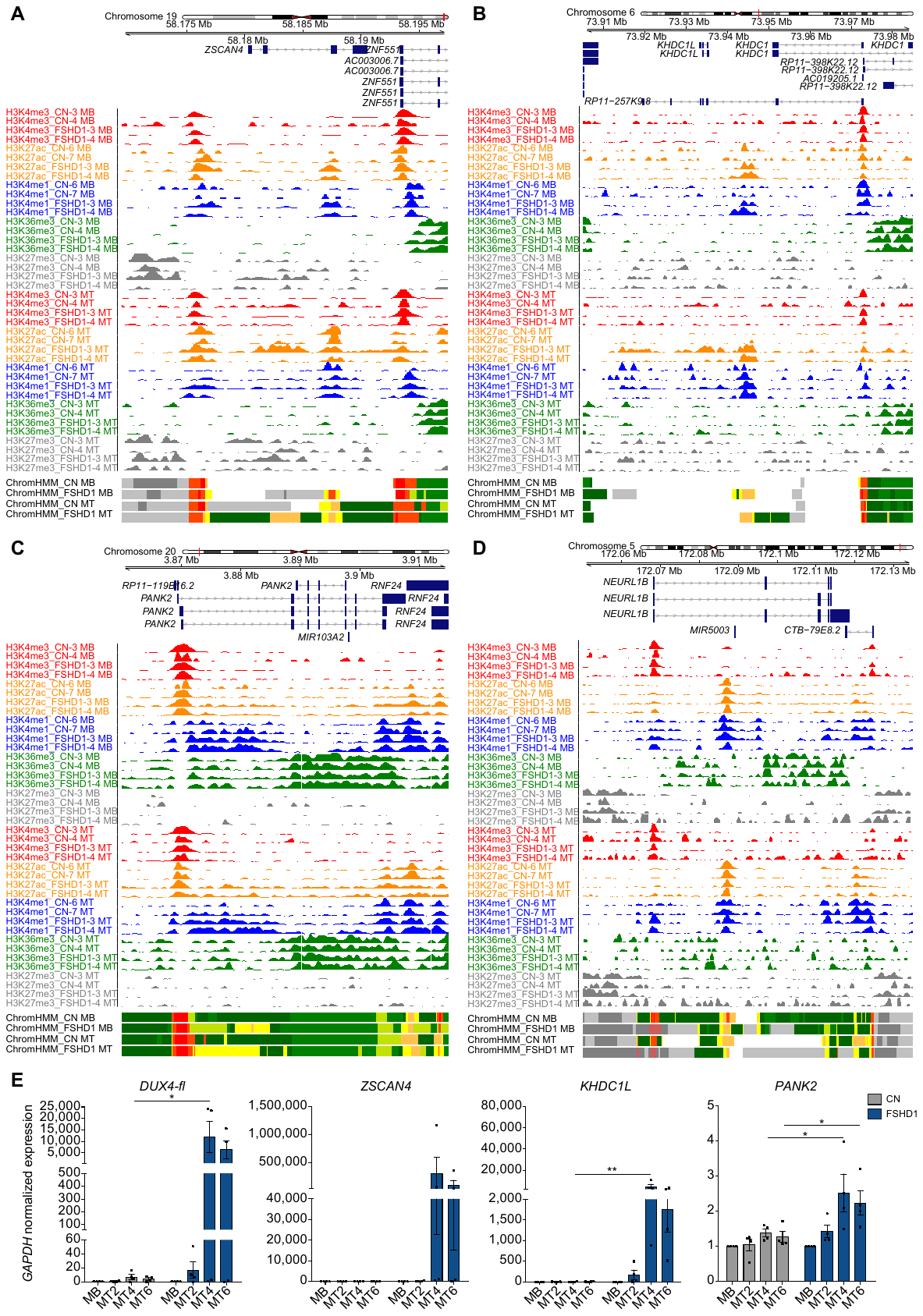

**Supplemental Figure S10. Examples of chromatin state definition of known DUX4 targets and their transcriptional levels**

(*A*-*C*) Representative ChIP-seq tracks of all histone modifications in each sample and chromatin states tracks of DUX4 upregulated genes *ZSCAN4*, *KHDC1L* and *PANK2* (Yao et al. 2014). (*D*) Representative ChIP-seq tracks of all histone modifications in each sample and chromatin states tracks of the DUX4 downregulated gene *NEURL1B* (Yao et al. 2014). (*E*) Expression levels of *DUX4-fl, ZSCAN4, KHDC1L and PANK2* genes during CN (grey) and FSHD1 (blue) differentiation (MB, myoblasts, MT2, myotubes day 2, MT4, myotubes day 4, MT6, myotubes day 6). Data were normalized on *GAPDH* expression and on MB. n=4 CN and 4 FSHD1. S.e.m. is indicated. *P* values were calculated by two-way ANOVA followed by Bonferroni post-test correction. Dots represent the values of each replicate; asterisks represent statistical *P* values; for *DUX4-fl* MT4, CN vs. FSHD1 *p*=0.0238; for *KHDC1L* MT4, CN vs. FSHD1 *p*=0.0015; for *PANK2* MT4, CN vs. FSHD1 *p*=0.0147, for *PANK2* MT6, CN vs. FSHD1 *p*=0.0440.

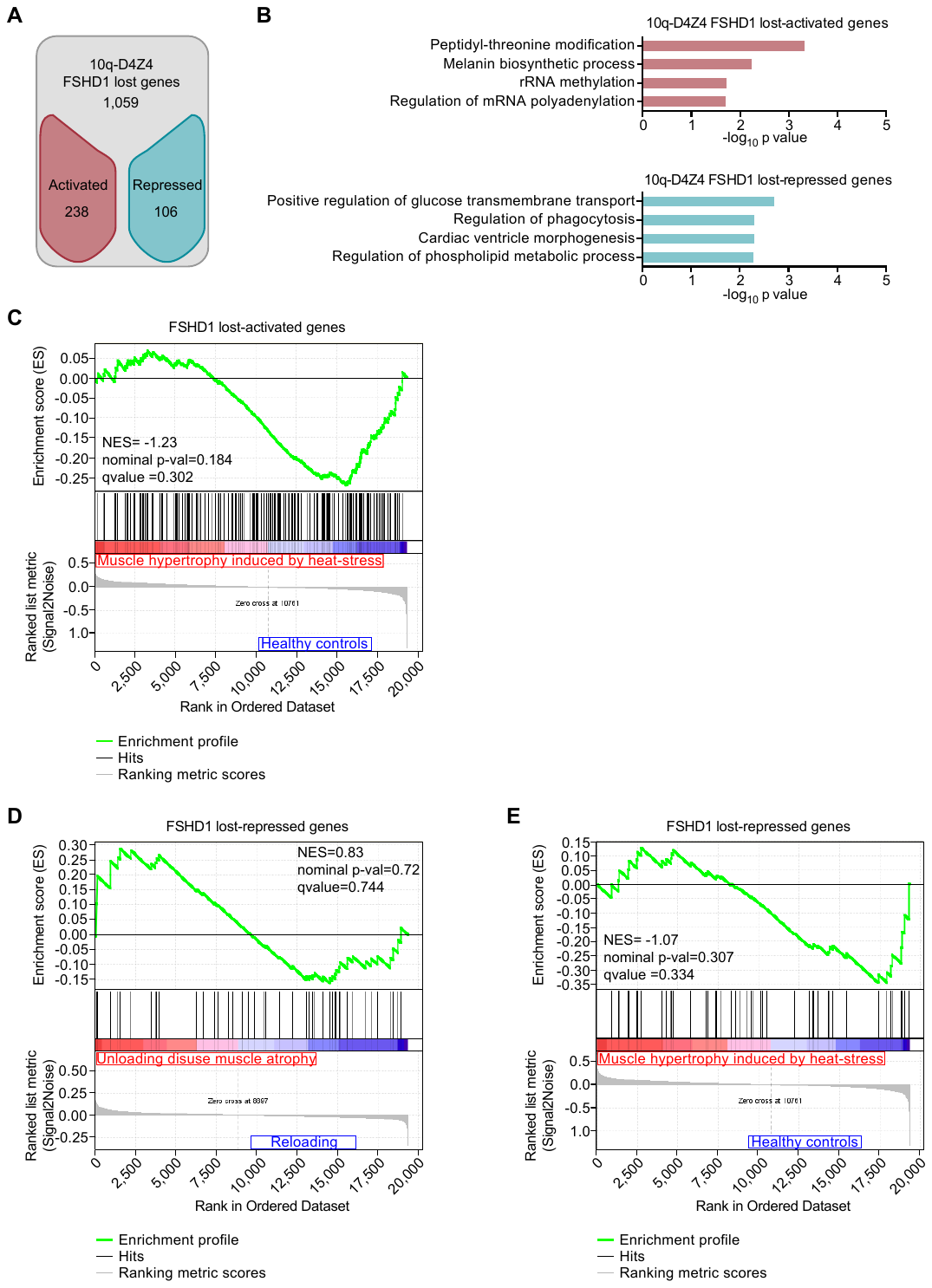

**Supplemental Figure S11. Additional data on 10q-D4Z4 FSHD1 lost genes and GSEA**

(*A*) Flowchart of filtering steps to identify 10q-D4Z4 FSHD1 lost genes. Genes within lost 10q-D4Z4-4C interactions were filtered as activated (red) or repressed (blue) in FSHD1. (*B*) Gene Ontology analysis (Biological Processes) of 10q-D4Z4 FSHD1 lost-activated and repressed genes. Bars correspond to -log_10_ of the *P* value. (*C*) Gene Set Enrichment Analysis (GSEA) results of the 319 FSHD1 lost-activated genes from Fig. 3A performed on expression data from heat-induced muscle hypertrophy (Goto et al. 2011). Genes upregulated in hypertrophy condition are depicted in red whereas genes not enriched are depicted in blue. NES, Normalized Enrichment Score. (*D*) GSEA results of the 131 FSHD1 lost-repressed genes from Fig. 3A performed on expression data from unloading-induced muscle atrophy subjects (Reich et al. 2010). Genes upregulated in atrophic condition are depicted in red whereas genes not enriched are depicted in blue. NES, Normalized Enrichment Score. (*E*) GSEA results of the 131 FSHD1 lost-repressed genes from Fig. 3A performed on expression data from heat-induced muscle hypertrophy (Goto et al. 2011). Genes upregulated in hypertrophy condition are depicted in red whereas genes not enriched are depicted in blue. NES, Normalized Enrichment Score.

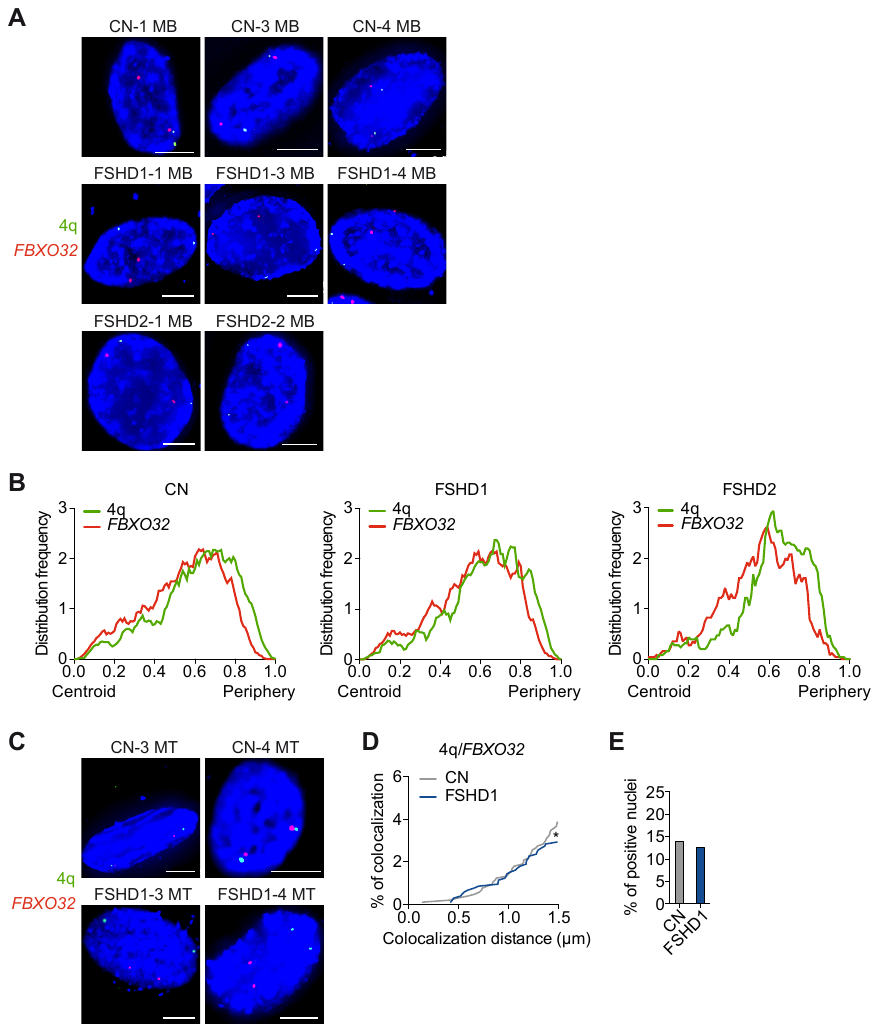

**Supplemental Figure S12. Additional information on *FBXO32/*4q-D4Z4 *trans* interaction**

(*A*) Representative nuclei of 3D multicolor DNA FISH using probes mapping to 4q35.1 region (4q, green) and *FBXO32* (red) in CN (CN-1, CN-3, CN-4), FSHD1 (FSHD1-1, FSHD1-3, FSHD1-4) and FSHD2 (FSHD2-1, FSHD2-2) myoblasts. Nuclei are counterstained with DAPI (blue). All images at 63X magnification. Scale bar=5 µm. (*B*) Frequency distributions of normalized distances from the nuclear centroid of 4q35.1 region (4q, green) and *FBXO32* (red) in CN (left), FSHD1 (middle) and FSHD2 (right) myoblasts. n= 1,826 (CN), 1,232 (FSHD1) and 510 (FSHD2). (*C*) Representative nuclei of 3D multicolor DNA FISH using probes mapping to 4q35.1 region (4q, green) and *FBXO32* (red) in CN (CN-3, CN-4) and FSHD1 (FSHD1-3, FSHD1-4) myotubes. Nuclei are counterstained with DAPI (blue). All images at 63X magnification. Scale bar=5 µm. (*D*) Cumulative frequency distribution of distances (below 1.5 µm) between 4q and *FBXO32* in CN (grey) and FSHD1 (blue) myotubes. n= 1,060 (CN) and 1,152 (FSHD1). *P* value was calculated by unpaired one-tailed *t*-test with confidence interval of 99%. Asterisks represent statistical *P* values; for CN vs. FSHD1 *p*=0.0283. (*E*) Percentage of nuclei positive for the interactions (under the cut-off of 1.5 µm). n= 265 (CN) and 288 (FSHD1).

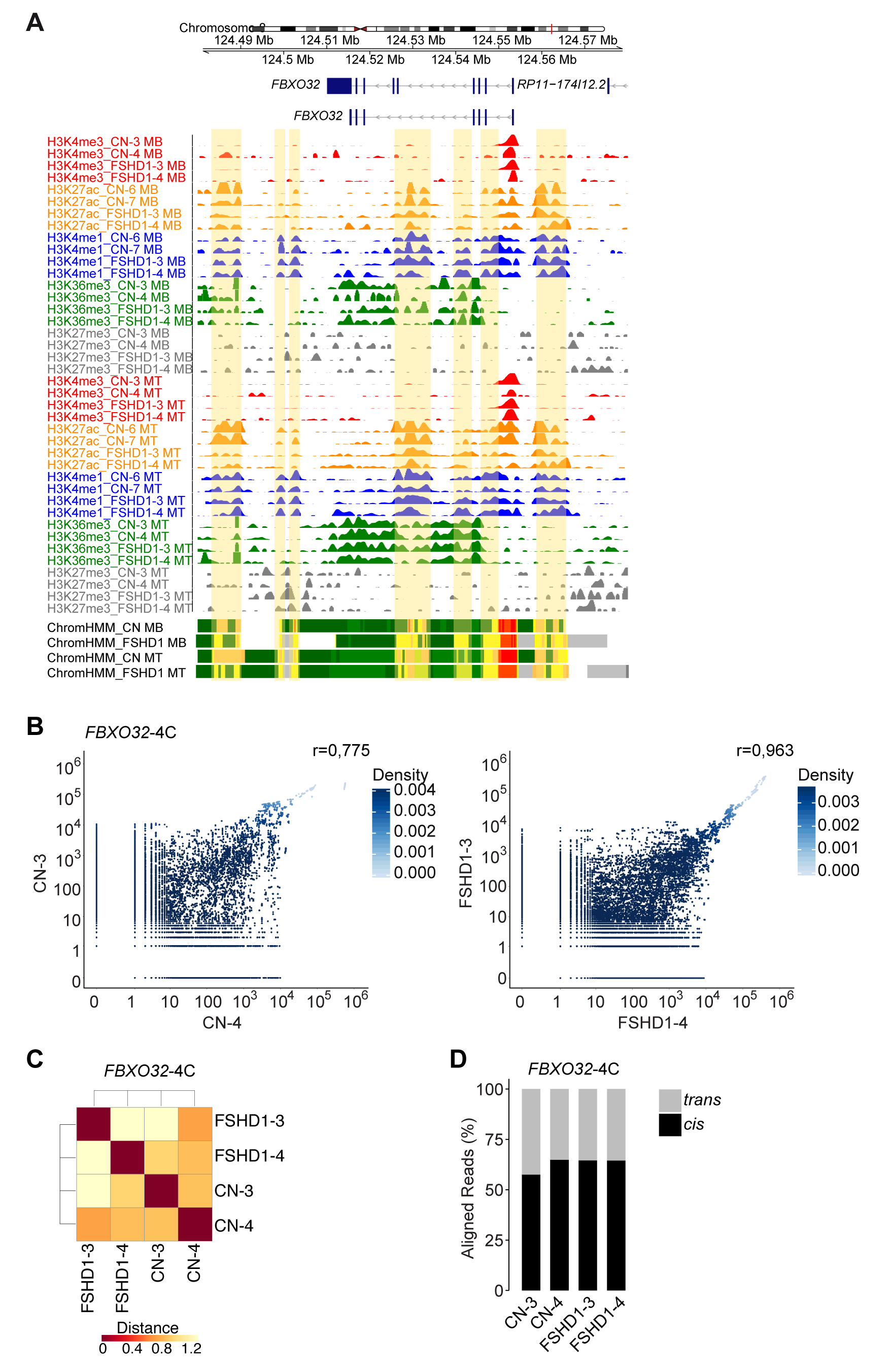

**Supplemental Figure S13. Additional information on FBXO32 4C-seq quality controls and *FBXO32* chromatin states**

(*A*) Representative ChIP-seq tracks of all histone modifications in each sample and ChromHMM chromatin states tracks at the *FBXO32* locus; enhancers are highlighted in yellow. (*B*) Scatter plots of running sum count reads for a window of 100 fragends for the *FBXO32*-4C. Pearson correlation coefficient shows the reproducibility between biological samples of CN (CN-3, CN-4) and FSHD1 (FSHD1-3, FSHD1-4). (*C*) Sample-to-sample comparison at the level of all fragends for the *FBXO32*-4C viewpoint. (*D*) Bar plot showing percentages of mapped reads *in* *cis* and *in* *trans* for the *FBXO32*-4C viewpoint.

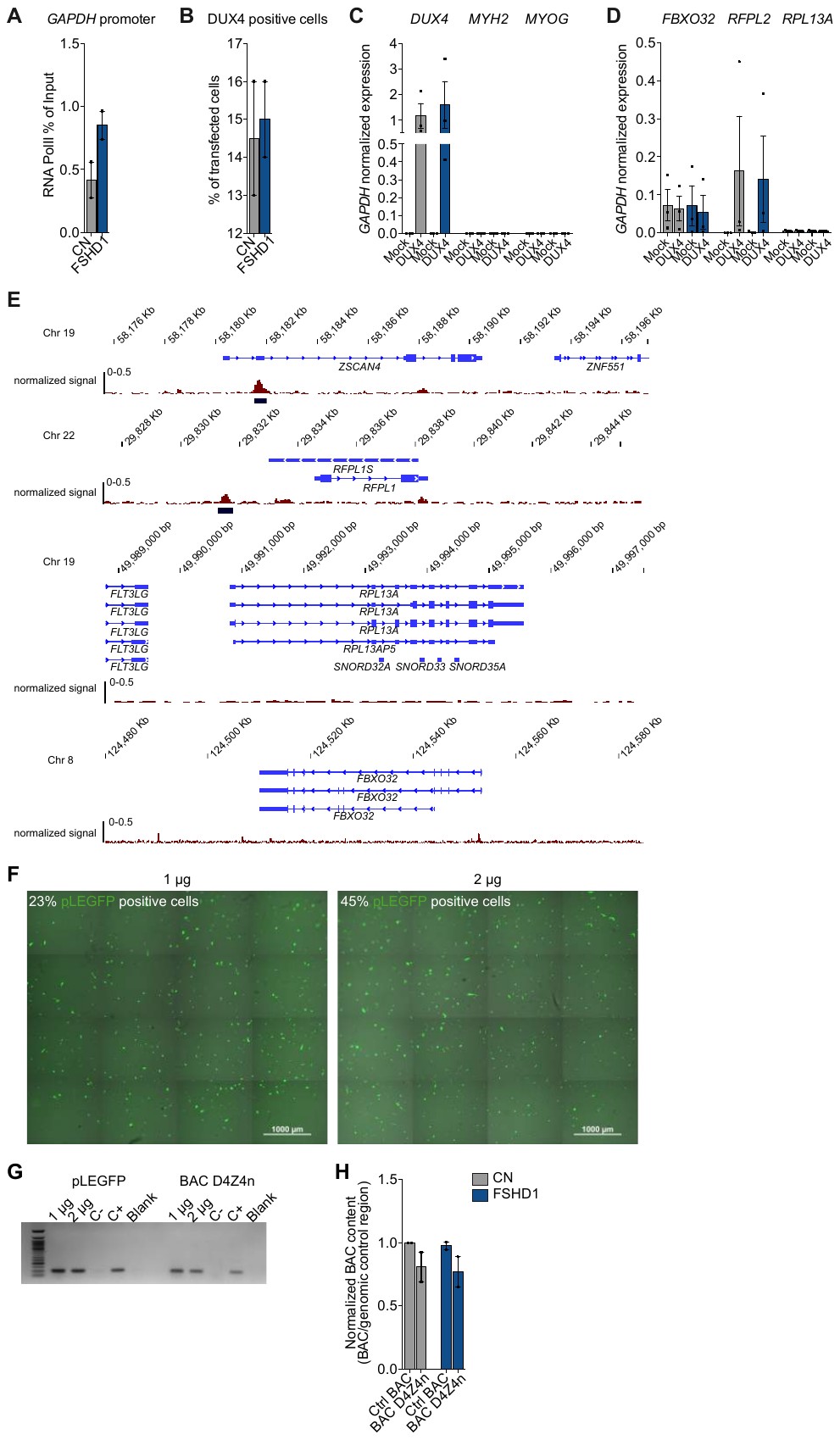
 **Supplemental Figure S14. Additional controls for *FBXO32* gene regulation and BAC transfection**

(*A*) Bar plots showing enrichment of RNA Pol II at *GAPDH* promoter assessed by ChIP-PCR experiment in CN (grey) and FSHD1 (blue) myoblasts. Results are presented as % of input. n=2 CN (CN-3, CN-4) and 2 FSHD1 (FSHD1-3, FSHD1-4). S.e.m. is indicated. Dots represent the values of each replicate. (*B*) Bar plot showing the percentage of cells transfected with a control plasmid (Mock) and a plasmid containing *DUX4* (DUX4) in CN (grey) and FSHD1 (blue) myoblasts. Percentages were calculated for 7,885 (CN) and 1,116 (FSHD1) counted nuclei. n=2 CN (CN-4, CN-5) and 2 FSHD1 (FSHD1-4, FSHD1-5). S.e.m. is indicated. (*C*) Bar plot showing expression levels of *DUX4*-ORF, *MYH2* and *MYOG* genes in CN (grey) and FSHD1 (blue) myoblasts transfected with a control plasmid (Mock) and a plasmid containing *DUX4* (DUX4). Data were normalized on *GAPDH* expression. n=3 CN (CN-3, CN-4, CN-5) and 3 FSHD1 (FSHD1-3, FSHD1-4, FSHD1-5). S.e.m. is indicated. Dots represent the values of each replicate. (*D*) Bar plot showing expression levels of *FBXO32* gene, *RFPL2* gene (DUX4 target; (Ferri et al. 2015)) and *RPL13A* gene (not a DUX4 target; (Ferri et al. 2015)) in CN (grey) and FSHD1 (blue) myoblasts transfected with a control plasmid (Mock) and a plasmid containing *DUX4* (DUX4). Data were normalized on *GAPDH* expression. n=4 CN and 4 FSHD1. S.e.m. is indicated*.* Dots represent the values of each replicate. *(E*) Genomic regions of genes positively and negatively regulated by DUX4 and DUX4 ChIP-seq peaks from published ChIP-seq dataset (Geng et al. 2012). (*F*) Light microscope images of human primary myoblasts transfected with 1 μg (left) and 2 μg (right) of pLEGFP (green). (*G*) Agarose gel run for PCR products of pLEGFP and BAC backbone regions on pLEGFP and BAC D4Z4n transfected (with 1 and 2 μg) myoblasts. (*H*) Bar plot showing normalized BAC content on a genomic control region assessed by PCR experiment in CN (grey) and FSHD1 (blue) myoblasts transfected with the control BAC (Ctrl BAC) and the BAC containing 4q upstream region and D4Z4 array (BAC 4q-D4Z4n). Data were normalized on the CN transfected with the control BAC. n=2 CN (CN-4, CN-5) and 2 FSHD1 (FSHD1-4, FSHD1-5). S.e.m. is indicated. Dots represent the values of each replicate.

**Supplemental Tables Legends**

**Supplemental Table S1.** Cell lines, SSLP and 4C-seq primers.

Sheet 1: Cell lines information

Sheet 2: SSLP information on cell lines analyzed in 4C experiments

Sheet 3: D4Z4-4C: primers and number of reads

Sheet 4: SSLP Read1 error rate assesments

Sheet 5: *FBXO32*-4C: primers and number of reads

**Supplemental Table S2.** 4C-seq interactions analysis.

Sheet 1: D4Z4 interactions summary

Sheet 2: Common and lost 4q-D4Z4 interactions in FSHD1

Sheet 3: Interactions of 4qA and 4qB alleles in CN and FSHD1

Sheet 4: 4qB vs 4qA lost interactions

Sheet 5: Common and lost 10q-D4Z4 interactions in FSHD1

Sheet 6: Specific and common interactions between 4q and 10q in CN and FSHD1

Sheet 7: 4q-D4Z4 vs 10q-D4Z4 lost interactions in FSHD1

Sheet 8: *FBXO32* nearbait interactions

**Supplemental Table S3.** 3D multicolor DNA FISH measurements.

Sheet 1: Colocalization distances

Sheet 2: Normalized distances from nuclear centroid

**Supplemental Table S4.** Chromatin state switches and RNA-seq expression levels.

Sheet 1: Chromatin state switches in MB

Sheet 2: Chromatin state switches in MT

Sheet 3: Expression levels of activated genes in MB

Sheet 4: Expression levels of repressed genes in MB

Sheet 5: Expression levels of activated genes in MT

Sheet 6: Expression levels of repressed genes in MT

**Supplemental Table S5.** Gene Ontology, GSEA and atrophic genes expression levels.

Sheet 1: Genes within 4q and 10q-D4Z4 lost interactions in FSHD1

Sheet 2: FSHD1 altered genes summary

Sheet 3: GO analysis for lost activated/repressed 4q-D4Z4 genes in FSHD1

Sheet 4: GO analysis for lost activated/repressed 10q-D4Z4 genes in FSHD1

Sheet 5: GSEA datasets

Sheet 6: Expression levels of atrophic genes in CN vs FSHD1 and FSHD2

**Supplemental Table S6.** Primers.

Sheet 1: 3C primers

Sheet 2: ChIP qPCR primers

Sheet 3: RT qPCR primers

Sheet 4: 3D FISH PCR probes
